# Haplotype Associated RNA Expression (HARE) Improves Prediction of Complex Traits in Maize

**DOI:** 10.1101/2021.04.30.442099

**Authors:** Anju Giri, Merritt Khaipho-Burch, Edward S. Buckler, Guillaume P. Ramstein

## Abstract

Genomic prediction typically relies on associations between single-site polymorphisms and traits of interest. This representation of genomic variability has been successful for prediction within populations. However, it usually cannot capture the complex effects due to combination of alleles in haplotypes. Therefore, accuracy across populations has usually been low. Here we present a novel and cost-effective method for imputing *cis* haplotype associated RNA expression (HARE, RNA expression of genes by haplotype), studied their transferability across tissues, and evaluated genomic prediction models within and across populations. HARE focuses on tightly linked *cis* acting causal variants in the immediate vicinity of the gene, while excluding *trans* effects from diffusion and metabolism, so it would be more transferrable across different tissues and populations. We showed that HARE estimates captured one-third of the variation in gene expression and were more transferable across diverse tissues than the measured transcript expression. HARE estimates were used in genomic prediction models evaluated within and across two diverse maize panels – a diverse association panel (Goodman Association panel) and a large half-sib panel (Nested Association Mapping panel) – for predicting 26 complex traits. HARE resulted in up to 15% higher prediction accuracy than control approaches that preserved haplotype structure, suggesting that HARE carried functional information in addition to information about haplotype structure. The largest increase was observed when the model was trained in the Nested Association Mapping panel and tested in the Goodman Association panel. Additionally, HARE yielded higher within-population prediction accuracy as compared to measured expression values. The accuracy achieved by measured expression was variable across tissues whereas accuracy using HARE was more stable across tissues. Therefore, imputing RNA expression of genes by haplotype is stable, cost-effective, and transferable across populations.

**Author summary:** The increasing availability of genomic data has been widely used in the prediction of many traits. However, genome wide prediction has been mostly carried out within populations and without explicit modeling of RNA or protein expression. In this study, we explored the prediction of field traits within and across populations using estimated RNA expression attributable to only the DNA sequence around a gene. We showed that the estimated RNA expression was more transferable than overall measured RNA expression. We improved prediction of field traits up to 15% using estimated gene expression as compared to observed expression or gene sequence alone. Overall, these findings indicate that structural and functional information in the gene sequence are highly transferable.

## Introduction

Genomic prediction is a powerful tool to predict quantitative traits using genomic information. In genomic prediction models, genome-wide predictors are simultaneously incorporated in the model in an attempt to capture variation from all quantitative trait loci (QTL) associated with the quantitative trait [1]. Genome-wide predictors could be single nucleotide polymorphisms (SNPs), haplotypes, or any downstream intermediate responses such as transcriptomes or metabolomes [1–6]. Haplotype sometimes yield higher prediction accuracy when compared to SNPs as they can capture local epistatic effects, can be in tight linkage with the QTL, and can better capture ancestral (identity by descent) relationships [7–10]. Haplotype-based models may be more useful as beneficial haplotypes are conserved across generations due to tight linkage. Downstream responses like gene expression may be biologically “closer” to the phenotype as they reflect transcription processes in different tissues. However, transcription is greatly affected by tissue, time, and growing conditions; therefore, transcriptome information from different tissues has varying power to predict phenotypes [2,4].

Gene expression is a complex phenomenon involving interaction between DNA, cell components, and the environment. Although every tissue in a plant contains the same genomic sequence, gene expression varies widely across tissues producing numerous phenotypes. The variation in gene expression is due to the differences in the regulatory regions and regulatory genes. Discerning the role of different factors contributing to expression is a challenge; however, a common approach to analyzing expression is to partition it into *cis* and *trans* components. The *cis* components are polymorphisms linked to the gene, whereas the *trans* are everything else not directly linked to the gene of interest [11]. *Trans* components are impacted by polymorphisms arising anywhere in the genome and affect gene expression by the products from diffusion and metabolism [12]. In maize like all eukaryotes, the expression of any gene is often impacted by dozens of transcription factors encoded in *trans* all across the genome [13]. Therefore, *trans* components frequently explain more variation in expression than *cis* components.

Different approaches exist to partition the variation in expression and infer the contribution to expression by *cis* factors only. These include hybrid crosses between inbreds and different testers to partition out background variation from *trans* [11,14], or analyses of genomic sequence linked to genes [15]. Here, we used haplotypes in the gene region and partitioned variation in expression contributed by the *cis* haplotype. Grundberg et al. [16] found that 90% of *cis* variants were shared across plants growing in different environmental conditions and only a few *cis* variants were environment specific as opposed to *trans* variants. The *cis* component of variation is less sensitive to genetic and environmental perturbation, so, they can be stable across different contexts and biological replicates. Partitioning out the variation due to *trans* from overall expression allowed us to get expression effect associated with the *cis* haplotype. We called this transferrable portion of the gene expression as the *cis H*aplotype Associated RNA Expression (HARE). We hypothesized that HARE would be more transferable across tissues than total measured transcript expression. Moreover, the consistent functional and structural information in HARE would result in higher prediction accuracy than total measured expression in predicting many complex traits.

We used maize to study transferability across different systems (tissues and populations) as it is an important cereal crop as well as an excellent model system for quantitative genetic studies [17]. Maize’s genotypic and phenotypic diversity has been explored in several studies using different mapping populations, uncovering thousands of genotypes and traits [18]. One example is the Goodman Association panel, which represents the global diversity of inbred genotypes from public maize breeding programs, including approximately 280 genotypes from tropical and temperate regions, sweet corn, and popcorn lines [19]. The nested association mapping panel (NAM) includes a set of approximately 5,000 recombinant inbred lines developed from 25 diverse inbreds crossed to a common parent, B73 [20,21]. NAM captures a large proportion of diversity in maize with less confounding by population structure as would occur in an association mapping panels. Both populations have been extensively genotyped and phenotyped for complex traits [22–25]. In addition, the Goodman Association panel also has a large set of available expression data from diverse tissues [26]. Recently these populations were used in the development of a Practical Haplotype Graph (PHG) utilizing high quality assemblies of NAM founder lines [27]. The PHG summarizes the diversity of these lines as a collection of haplotypes in a graph [27]. In diverse species like maize, with rich allelic series, a wide range of possible alleles might result in the same molecular outcome (e.g., gene expression, protein expression, etc.), a process known as equifinality. HARE can parsimoniously summarize a large series of allelic variants including causal variants, resulting in transferability across populations. Therefore, we hypothesized that HARE would be functionally relevant beyond genomic relationships and would result in higher prediction accuracy than haplotype structure when used to predict many complex traits within and across populations.

To test these hypothesis, we designed a novel method for imputing expression associated with haplotypes in the genic regions by HARE and studied the transferability of imputed expression across tissues and populations. The HARE estimates were imputed in NAM founder’s genic haplotypes using gene expression data previously collected in seven diverse tissues [26]. The objectives of the study were to: i) partition gene expression variation into *cis* and *trans* components, ii) impute HARE in NAM and the Goodman Association panels based on the shared NAM founder’s haplotypes, iii) assess prediction of many complex traits by using HARE, randomly permuted HARE (preserving haplotype structure only), and measured expression within and across populations, and iv) integrate HARE from different tissues to predict complex phenotypes within and across populations.

## Results

### Phenotypic and genetic diversity in NAM and Goodman Association panel

The phenotypic distribution of 26 diverse traits is presented in S1 Fig, where the average trait value was higher in 18 traits in NAM than Goodman Association panel. The haplotype frequency was also variable across these two panels (S2 Fig). The median haplotype count across genes (reference regions) was 100 in NAM whereas it was 8 in the Goodman panel. The majority of haplotypes were present in 100 lines as expected from biparental populations with 200 inbred progenies in NAM. Each reference region in NAM was dominated by haplotypes from the common parent (B73), representing half of the haplotypes. We also calculated haplotype entropy from haplotype frequency counts in each reference region to reflect the average information content of haplotypes in reference regions. We observed a higher median entropy of 3.03 in Goodman panel when compared to 2.3 in NAM (S2 Fig).

### Variance partition in expression

We hypothesized that the majority of the expression would be contributed by *trans* acting factors as compared to the *cis* component. To test this hypothesis, we fit the model with haplotypes in each gene region as *cis* and the haplotype relationship matrix (HRM) combined across all genes as *trans* (model 1; Fig 1, B). For most of the genes, higher variance in RNA expression was explained by the *trans* component (Fig 2a right) as compared to the *cis* component (Fig 2a left), irrespective of the tissue. Overall, *cis* haplotypes contributed only 34% (31-38% across individual tissues) of the total genetic variation in expression across all tissues.

**Fig 1.**
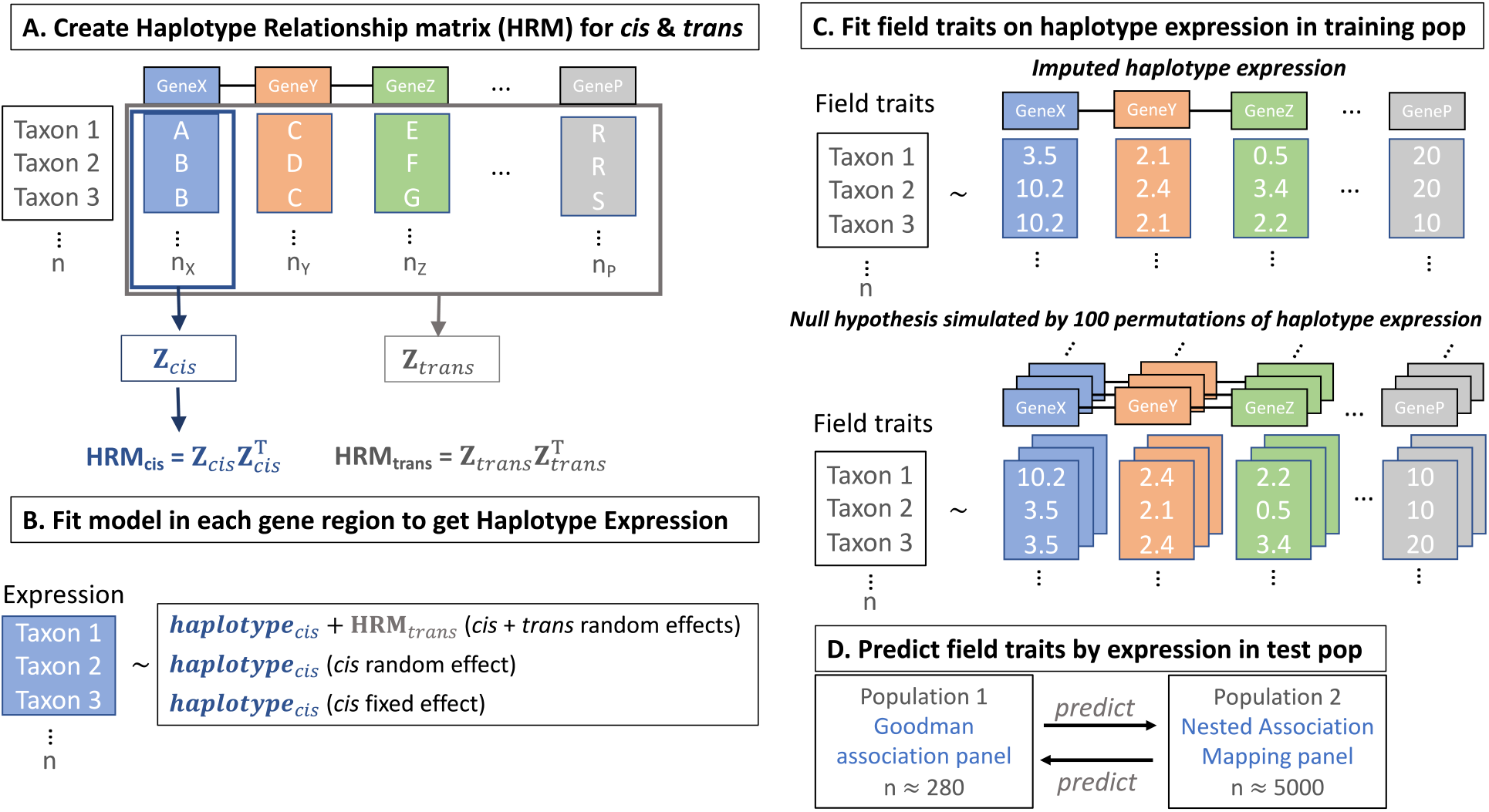
Experimental methods on calculating haplotype associated RNA expression (HARE) and using HARE values to predict complex traits. A. The haplotypes of 26 NAM founders and one additional stiff stalk inbred line were identified in each gene region of the Goodman Association panel by mapping GBS reads (presented in detail in Valdes Franco et al. [27]) to the indexed pangenome of 27 lines. Haplotype relationship matrices (HRM_*cis*_) were created in each gene region and all genic HRM_*cis*_ were combined to get HRM_*trans*_ to control for *trans* effects, B. Using gene expression from seven tissues [26], fixed or random effects models were fitted in each gene region with or without controlling for *trans* effects for each gene, C. Models were trained using field phenotypes in the Goodman panel or the NAM panel using HARE estimates or 100 randomly permuted HARE values while preserving haplotype structure, and D. Trained models were used to predict complex traits across populations.

The total heritability was quantified as the proportion of variation contributed by *cis* and *trans* components to the total variation in gene expression. Overall, the gene expression was highly heritable with an average ranging from 50% to 59% across tissues (Fig 2b right). Though the gene expression was highly heritable, the heritability was largely contributed by *trans* compared to *cis* (Fig 2b). The large effect of *trans* could be due to many small-effect molecular connections from *trans* regulators [11,28].

**Fig 2.**
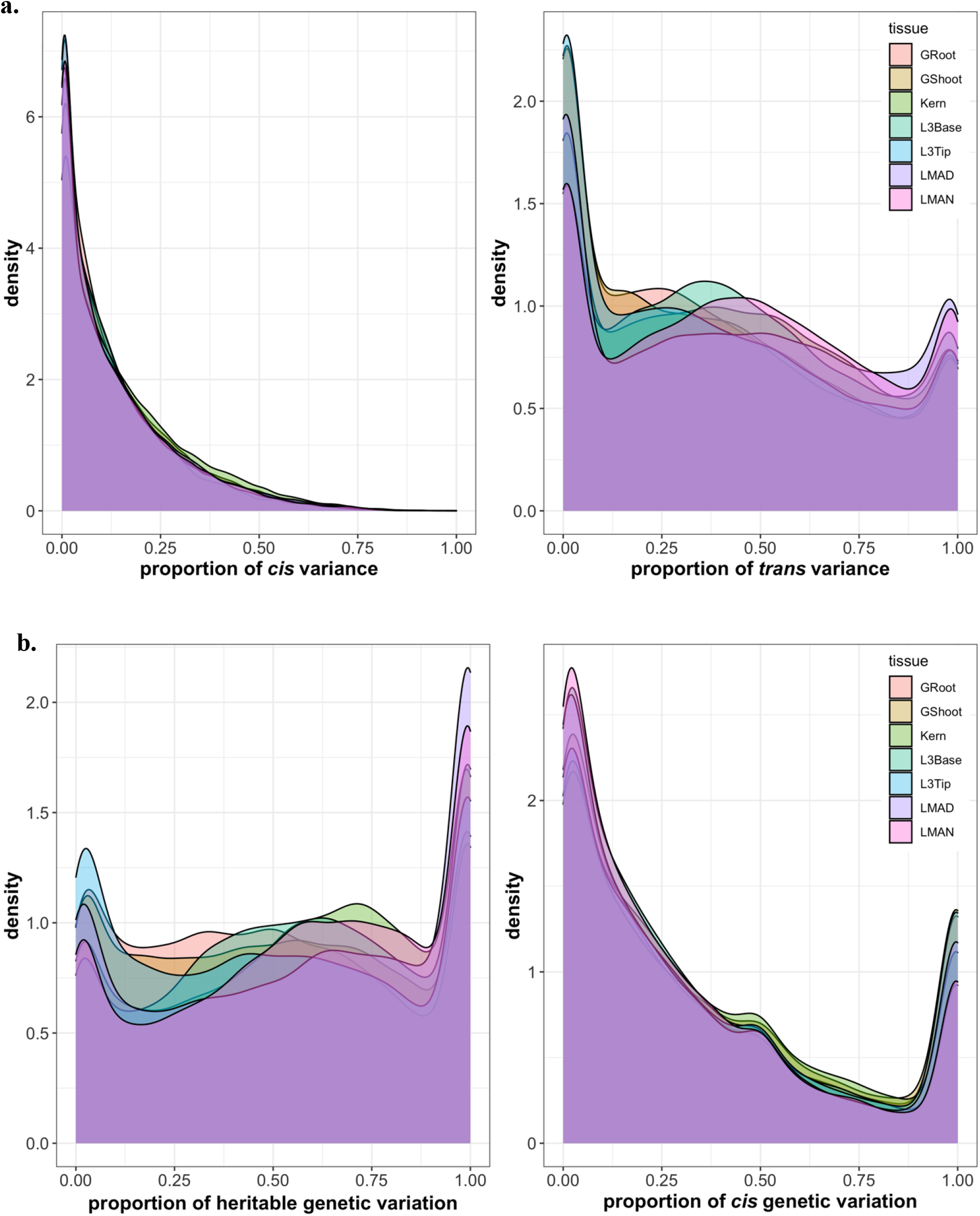
*Cis* haplotype explained one-third of the total genetic variance in expression. a. Proportion of variation explained by *cis* (left) and *trans* (right) components, b. Proportion of heritable genetic variation over phenotypic variation (left) and proportion of heritable genetic variation explained by *cis* (right) in gene expression across seven different tissues. Heritable variation was calculated in each gene as the ratio of the sum of *cis* and *trans* variance to total variance. Different colors represent seven diverse tissues in maize: germinating seedlings root (GRoot), germinating seedlings shoot (GShoot), two cm from base of leaf 3 (L3Base), two cm from tip of leaf 3 (L3Tip), mature mid-leaf tissue sampled during mid-day (LMAD), mature mid-leaf tissue sampled during mid-night (LMAN), and developing kernels harvested after 350 growing degree days after pollination (Kern).

We analyzed the result separately for a set of genes (~8,000 genes) with higher expression across each tissue to see if the patterns in their expression were any different from other genes. The *cis* haplotype explained a similar amount of variation (median 33%), however, the heritability increased slightly from a median of 54% to 60% across all tissues in a set of highly expressed genes.

### Transferability of Haplotype Associated RNA expression (HARE)

We used three different models to estimate HARE. Model 1 included both *cis* and *trans* fit as random effects, and model 2 and model 3 included only *cis* fit either as a fixed or random effect. To determine how close the HARE estimates were to measured expression, we estimated their correlations across all tissues. A correlation close to 1 implied that the majority of variation in gene expression was contributed by *cis*, whereas a correlation close to 0 meant most expression was contributed by *trans*. The overall distribution across all expressed genes was similar in all tissues and models with a mean correlation of 0.44 (Fig 3).

**Fig 3.**
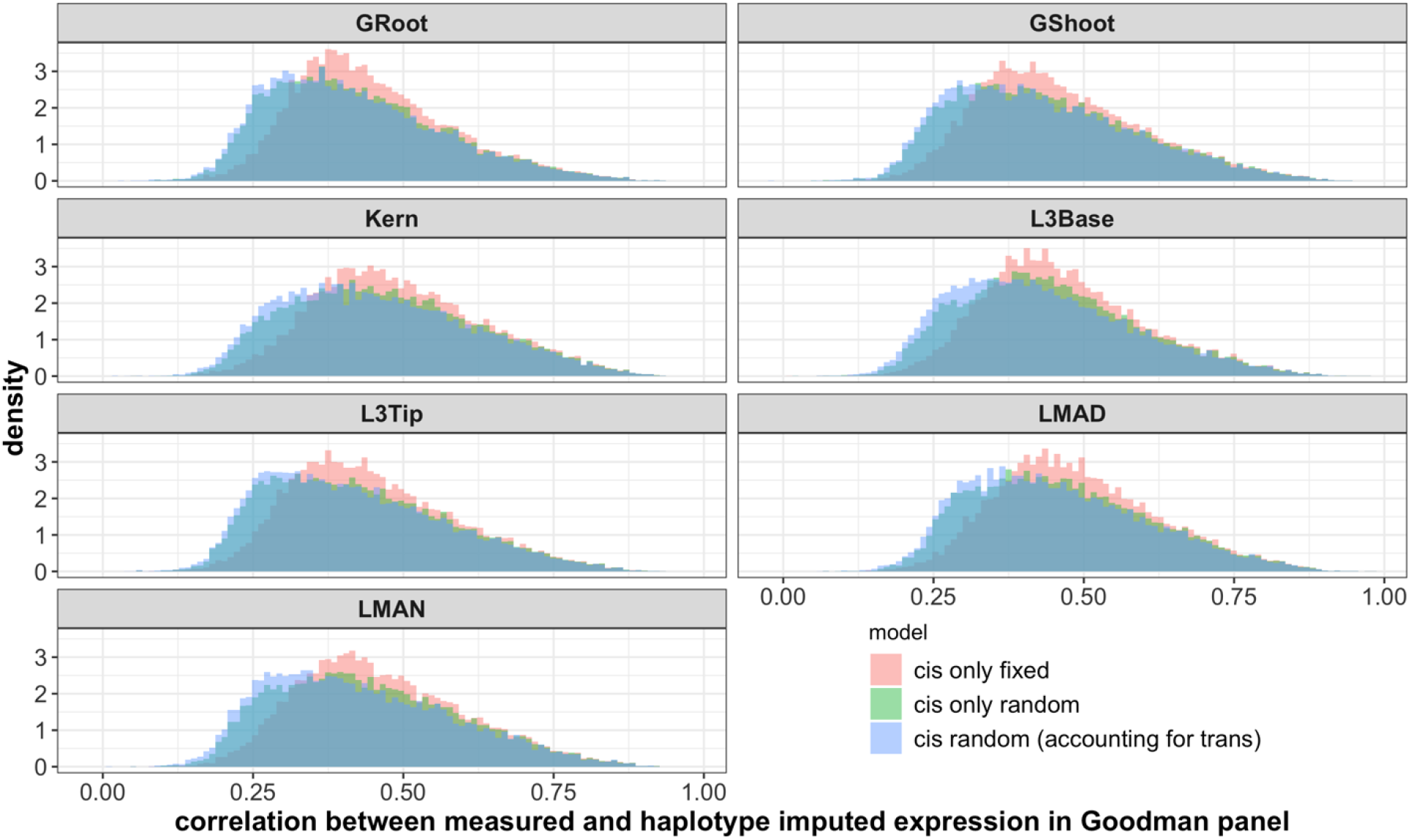
Haplotype associated RNA expression (HARE) was moderately correlated with measured RNA expression across seven diverse tissues. The different colors represent HARE imputed from three statistical models: Model 1 (*cis* fixed effect), 2 (*cis* random effect), and 3 (*cis* + *trans* random effects). Transcripts from measured RNA expression was a result of genetic signals in both *trans* and *cis*. Therefore, the correlation was moderate for most of the genes.

We hypothesized that HARE would include the transferable portion of gene expression based on the underlying haplotype. To test this, we looked at pairwise correlations of HARE from all three models and measured transcript expression across all combinations of genes and tissues. Correlation coefficients ranged from −1 to 1 in all 21 different combinations of 7 diverse tissues (Fig 4, S3 Fig). The median correlation coefficient in measured expression was 0.14 across all tissues, whereas it was 0.4 in HARE. Correlation across tissues was larger for a set of highly expressed genes (~8,000 genes) as compared to the overall set. The median correlation across tissues increased from 0.14 to 0.21 in measured expression and 0.4 to 0.53 in HARE (S5 Fig) in the highly expressed gene set. The HARE imputed from all three models followed similar trends of higher correlation for most of the genes across all tissues as compared to measured transcript expression. Closely related tissues -- for example, mature mid-leaf tissue sampled during midday (LMAD) and midnight (LMAN) were more correlated than other combinations, both in measured expression and HARE (S3 Fig), reflecting the influence of shared gene regulation mechanisms driving these correlations in leaf tissues.

**Fig 4.**
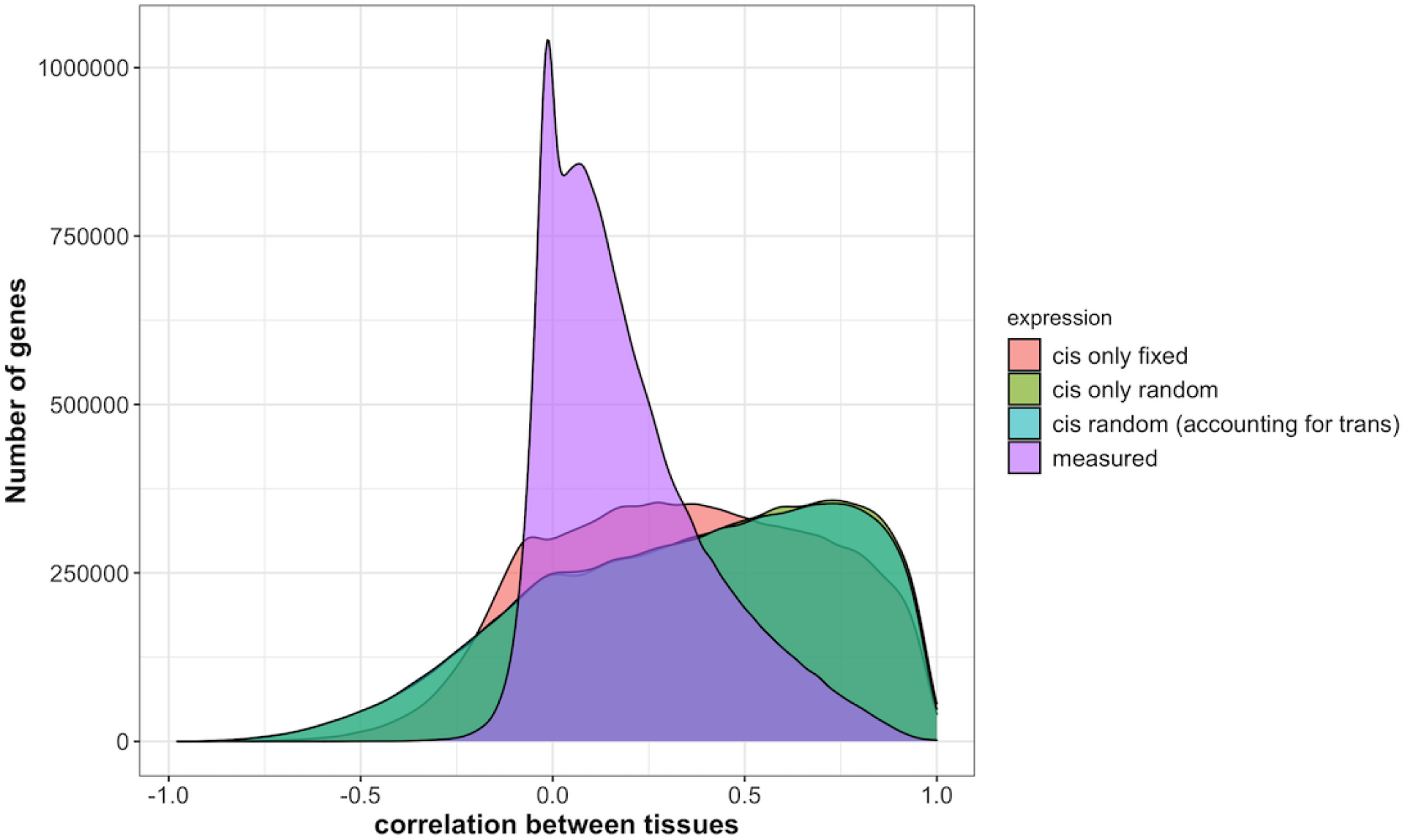
Haplotype associated RNA expression (HARE) was highly correlated across tissues as compared to measured transcript expression. Different colors represent HARE imputed from 3 statistical models: Model 1 (*cis* fixed effect), 2 (*cis* random effect), and 3 (*cis* + *trans* random effects), and measured transcript expression. The distribution is pairwise correlation of genes across 21 different combinations from 7 different tissues.

We looked further into highly correlated genes (correlation > 0.75) and poorly correlated genes (correlation −0.05 to 0.05) across all tissue combinations to see if the large correlations were the result of some very lowly expressed genes. We examined expression in each tissue and filtered values for genes with fragments per million counts <5. In tissue LMAN, out of 10,600 genes with low correlation across tissues, only 3,000 lowly expressed genes were filtered out. Out of 11,500 genes with high correlation across tissues, only 1,400 genes showed low expression values across tissues. Therefore, the lowly expressed genes did not drive the higher correlation of HARE estimates across tissues.

### Genomic prediction using HARE

The high correlation of HARE estimates across tissues suggests that consistent and transferable genetic information is present in HARE. The functional application of HARE was evaluated using genomic prediction within and across populations for 26 agronomically important traits in maize (Table 1). Transferability across populations was evaluated based on prediction accuracy, calculated as the Pearson correlation of observed and predicted trait values. The genomic prediction models were trained to predict traits within and across populations in maize. First, we compared prediction accuracies in three sample traits (days to anthesis, days to silking, and plant height) using HARE estimates and their randomly permuted values (“random HARE”, representing only haplotype structure; see Materials and methods), from three different methods (models 1, 2, and 3). We did not see any significant differences in accuracy using any of these imputation methods (S6 Fig), so we used HARE estimates from model 1 (*cis* effects adjusted from *trans* effects) to predict all 26 traits within panel in the Goodman Association panel and across panels in the Goodman Association and NAM.

**Table 1.** Selected traits for genomic prediction

| Category | Traits | Reference |
| --- | --- | --- |
| **Flowering** | Days to silking | [25] |
|  | Days to anthesis | [25] |
|  | Anthesis silking interval | [25] |
|  | Tassel length | [22] |
|  | Tassel primary branches | [22] |
| **Morphology** | Plant height | [25] |
|  | Ear height | [25] |
|  | Leaf length | [22] |
|  | Leaf width | [22] |
|  | Leaf angle | [22] |
|  | Nodes below ear | [22] |
|  | Nodes above ear | [22] |
|  | Number of brace roots | [22] |
| **Yield related** | Cob diameter | [22] |
|  | Cob length | [22] |
|  | Ear row number | [22] |
|  | Kernel number per row | [22] |
|  | Ear mass | [22] |
|  | Cob mass | [22] |
|  | Kernel wt | [22] |
|  | Test wt | [22] |
|  | Total kernel number | [22] |
| **Kernel composition** | Starch | [24] |
|  | Protein | [24] |
|  | Oil | [24] |
| **Disease** | Southern leaf blight | [23] |

### Within-panel prediction using HARE as compared to measured expression or haplotype structure (random HARE)

The comparison of prediction accuracy using measured expression and HARE was conducted in the Goodman panel to test the hypothesis that the prediction using HARE would be higher than measured expression. Prediction accuracy by measured expression was highly variable across traits and tissues; in contrast, prediction accuracy by HARE was less variable across tissues. The highest accuracy was observed for flowering time traits (e.g., days to anthesis up to 0.9), using HARE from all tissues, or measured expression from mature mid-leaf tissues (LMAD and LMAN). Overall, HARE resulted in higher prediction accuracy for *all* 26 traits, compared to measured expression in any tissue (Fig 5a and S7 Fig). The highest accuracy increase was for the number of brace roots, which increased from 0.21 to 0.5 using HARE from germinating root (S7 Fig). However, the median increase across tissues was highest for trait kernel weight, which increased from 0.2 to 0.5 (Fig 5a). Also, HARE resulted in significantly higher prediction accuracy compared to random HARE for 24 traits (P-value < 0.05) (Fig 5b). Therefore, partitioning expression at the level of gene haplotypes results in higher prediction accuracy, when compared to predictions by measured expression or haplotype structure (random HARE).

**Fig 5.**
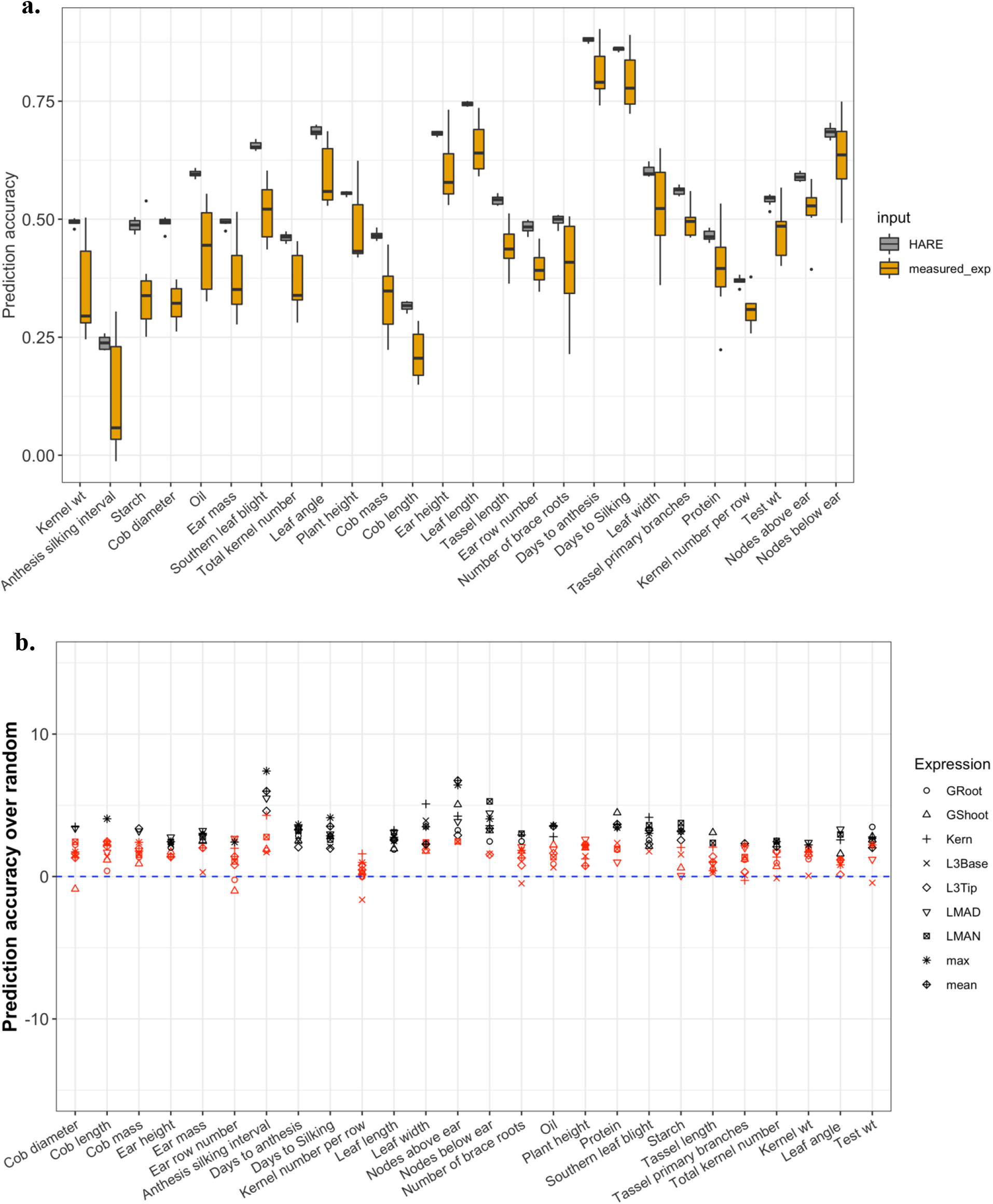
HARE improved within-panel prediction accuracy over measured expression and random HARE for most of the traits. (a) Prediction accuracy within the Goodman Association panel using HARE and measured expression (measured_exp) from all tissues arranged by prediction differential. (b) Change in prediction accuracy using HARE over the mean accuracy from random HARE (blue dashed line). Different symbols represent HARE from different tissues. The black shapes represent statistically significant differences at P-value <0.05 and red shapes are without significant differences. P-values calculated using Monte Carlo procedure.

### Cross-panel prediction using HARE as compared to haplotype structure (random HARE)

For all 26 traits and 7 diverse tissues, prediction models were trained using HARE or random HARE across panels in NAM and the Goodman Association to determine if HARE carries functional information beyond haplotype structure. HARE often improved prediction accuracy of many traits when the model was trained in NAM or the Goodman panel as compared to random HARE. HARE significantly increased accuracy by 34.6% and decreased 1.7% across all traits and tissue combinations when the model was trained in NAM (Fig 6a and S1 Table) and only significantly increased by 21.8% when trained in Goodman (Fig 6b and S2 Table). The accuracy was significantly higher in 17 out of 26 traits when trained in NAM and tested in the Goodman panel, *versus* 19 out of 26 traits when trained in Goodman panel and tested in NAM (P-value < 0.05). However, the increase in accuracy was highly variable across these two panels. The increase was as high as 15% (for morphological traits: plant height and leaf length) when the model was trained in NAM and tested in the Goodman panel; whereas it was less than 10% when the model was trained in the Goodman panel and tested in NAM (Fig 6a, 6b, and 7). The increase in accuracy over random HARE was also observed when a model was trained only in the sample of 250 NAM RILs (a similar size as the Goodman panel, 10 sample RILs from 25 families) to predict 3 traits (Days to anthesis, Days to silking, and Plant height) in Goodman panel (S8 Fig). When the model was trained in NAM the increase in prediction accuracy reached up to 16% over random HARE for the morphological traits, 10% for flowering traits, 8% for yield traits, 12% for kernel composition, and 6% for disease related traits (S1 Table). In general, traits in yield and disease related categories had the lowest accuracy when compared to the traits in other categories. Genomic prediction models using HARE could improve prediction accuracy with simple computational work without any additional cost for data generation. Therefore, haplotype-based models have the potential to improve genomic prediction across populations; however, the improvement depends on the traits of interest. The overall number of significant improvements was higher when using mean or maximum expression, when compared to using the expression of individual tissues, in both directions. Therefore, integrating expression from diverse tissues as mean or maximum expression may contribute to further improvements in prediction accuracy.

**Fig 6.**
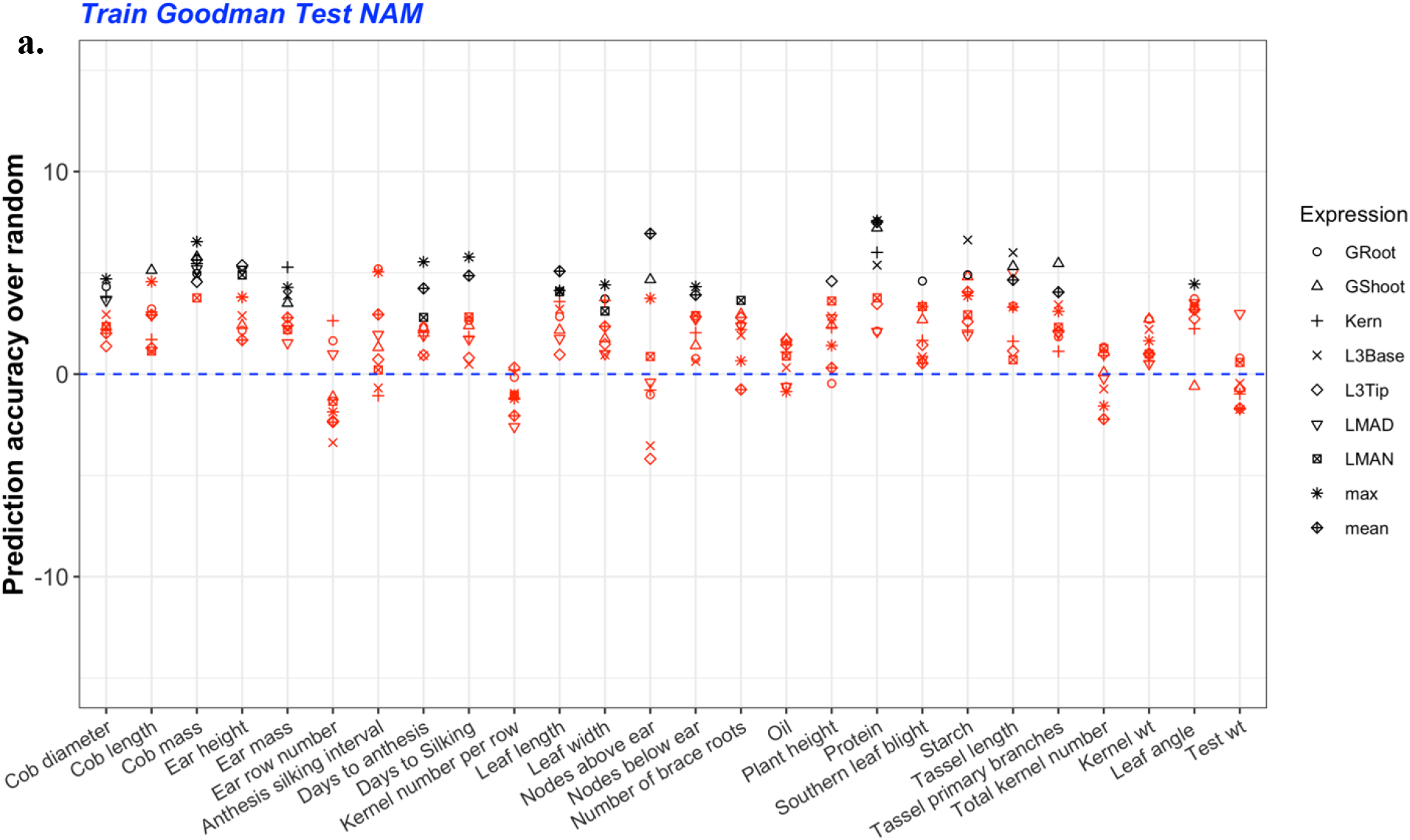
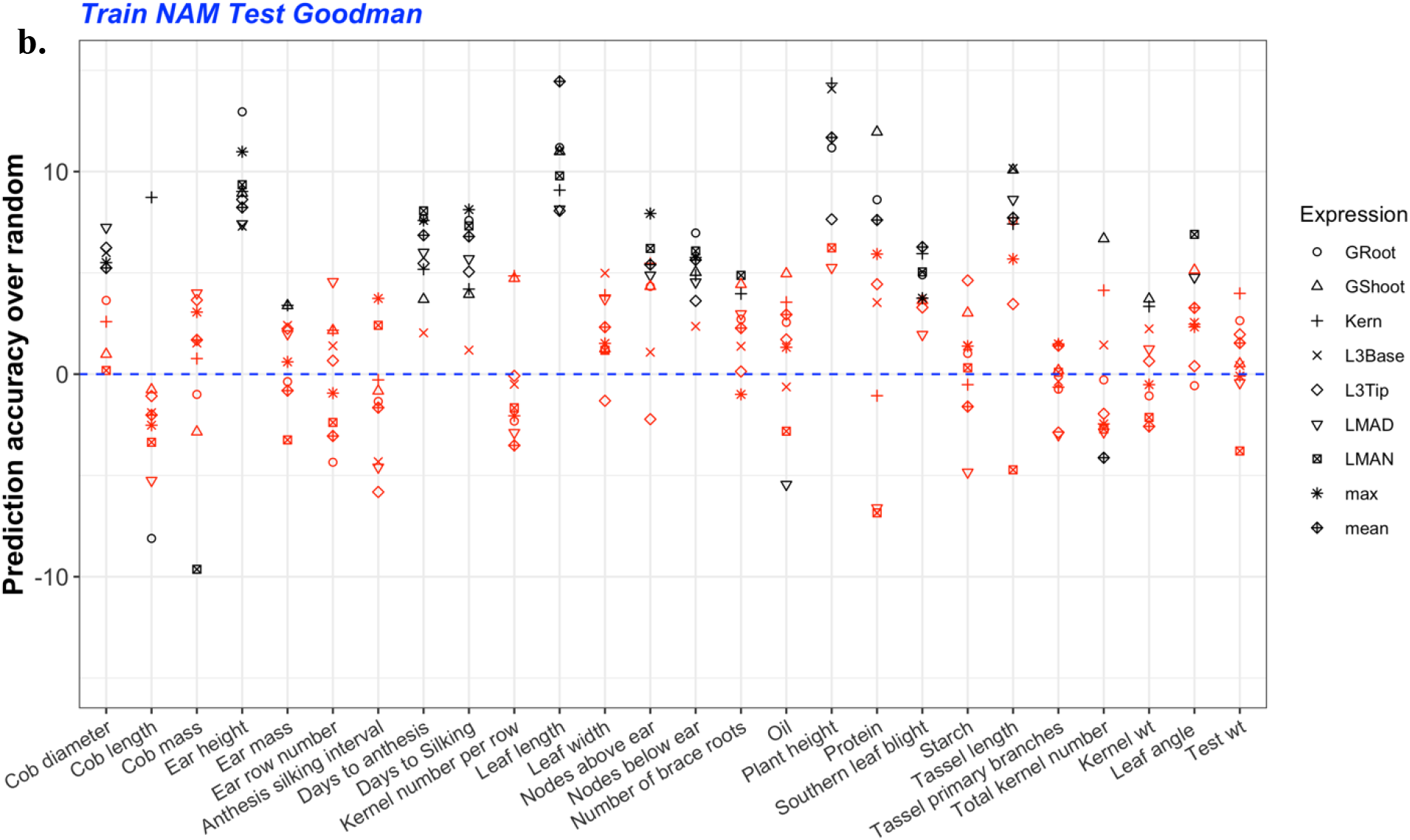
HARE improved cross-panel prediction accuracy over random expression values for most of the traits. Change in prediction accuracy using HARE over the mean accuracy from random HARE (blue dashed line) for models (a) trained in the Goodman panel and tested in NAM, (b) trained in NAM and tested in the Goodman panel. The different symbols represent HARE from different tissues: germinating seedlings root (GRoot), germinating seedlings shoot (GShoot), two cm from base of leaf 3 (L3Base), two cm from tip of leaf 3 (L3Tip), mature midleaf tissue sampled during mid-day (LMAD), mature mid-leaf tissue sampled during mid-night (LMAN), and developing kernels harvested after 350 growing degree days after pollination (Kern). The black shapes represent statistically significant differences at P-values <0.05 and red shapes represent no statistical significance. P-values were calculated using a Monte Carlo procedure.

**Fig 7.**
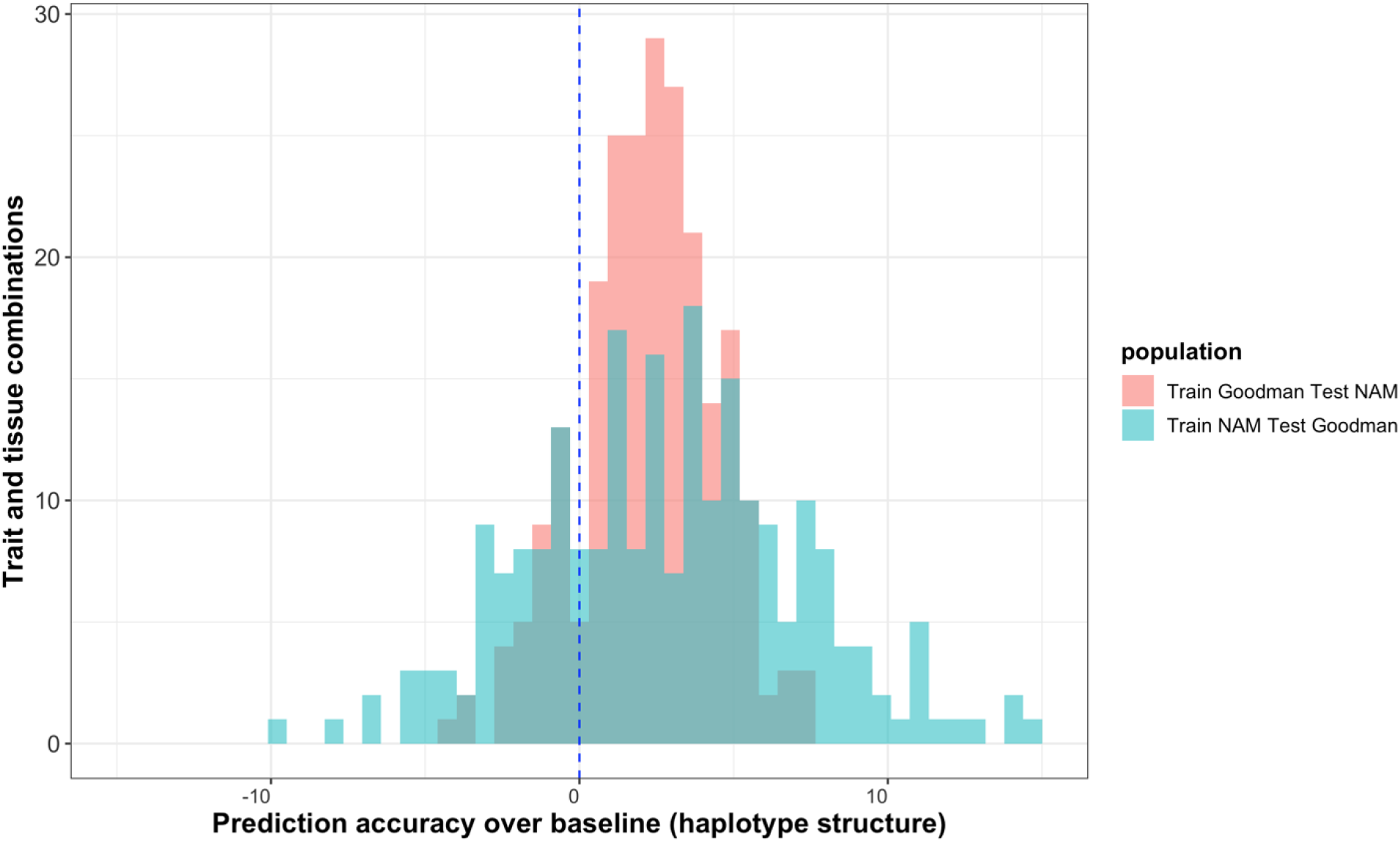
HARE increased prediction accuracy by up to 14% when the model was trained in NAM and tested in the Goodman panel. Summarized differences in prediction accuracy using HARE over the mean accuracy from random HARE (representing haplotype structure) across 26 phenotypes and 7 diverse tissues. The blue dashed line is the mean of the average prediction accuracy using random HARE across each trait and tissue combination.

## Discussion

### Cis haplotypes explained one-third of the genetic variation in expression

Consistent with other studies, we found the vast majority of expression to be heritable [28–30]. In eQTL mapping, *cis*-eQTL can seem predominant as they are frequently the single largest QTL for a given gene, but this is likely a power and multiple testing issue [12]. By using variance partitioning and assuming a polygenic model, we are likely accurately estimating the relative importance of these two components, *cis* and *trans*.

We showed that *cis* haplotype explained around 34% of variation in expression, which given the rapid linkage disequilibrium decay surrounding the gene is still extremely enriched for variance. Similar results were observed by Lemmon et al. [30] in maize and teosinte using hybrid allele specific expression, a complementary technique to our approach. This agree with our molecular knowledge, where, dozens of transcription factors likely regulate each gene [13]. These transcription factors are a result of any regulatory genes modelled as *trans*. In contrast, *cis* variability is a result of variation within or around genes only, empirically lowering the amount of variability explained by a *cis* as compared to the overall *trans* effect as observed in similar experiments in human and yeast [12,29].

### HARE was highly transferable across tissues as compared to measured transcript expression

Variation in gene expression across tissues, developmental stages, genotypes, and experimental conditions has been shown in earlier studies in plants and humans [26,33–35]. Lower correlations in expression for similar genes have been observed across populations in Mogil et al. [36], therefore, high gene expression in one population may not always be as high in diverse panels. The lack of strong correlations in measured transcript expression may result from *trans* effects in the expression, which are specific to tissues, genetic backgrounds, or environmental conditions [11] (S4 Fig). With HARE, we observed higher transferability across tissues as this portion of the variation in gene expression was less sensitive to environmental perturbations and biological contexts. The *cis* regulatory mutations affect expression of fewer genes compared to *trans* effects, so, they result in less pleiotropy and involve fewer functional tradeoffs [37]. In the absence of large pleiotropic effects, selection can act more consistently, so *cis* effects may be more transferable across different backgrounds [37,38].

HARE can integrate a rich allelic series that is more transferable across different contexts than measured transcript expression which is a result of *cis*, *trans*, their interaction, and environmental effects. Allelic richness is more pronounced in species like maize, which has 20 times higher nucleotide diversity than human beings [39]. Because of high allelic richness in maize, a wide range of possible alleles might lead to the same molecular outcome (for example, gene expression, protein expression, etc.), a concept known as equifinality. Due to equifinality, it has been observed that allelic variants are not always shared across genomes, and transcription is not always correlated with translation [40]. However, the *cis* portion of expression that summarizes allelic richness is highly transferable across tissues. The *cis* variants in HARE located in the close promoter, 5’ and 3’ untranslated regions, introns and the gene regions are likely consistent across tissues. Further research is needed to understand the effect of enhancer and tissues interaction in the variation on *cis* effects, however the current study suggests that it’s not the dominant factor.

### HARE improved prediction over transcriptome expression

Biological information flows along the central dogma from the genome to transcriptome to proteome to metabolome, and finally to complex phenotypes [41]. For most trait and tissue combinations, transcriptome expression yielded lower prediction accuracy when compared to HARE (Fig 5a and S7 Fig). Furthermore, we observed less variability in prediction accuracy using HARE, which points to the context dependence of RNA expression. HARE owes its consistent advantage in prediction accuracy to functional information that does not include non-genetic sources of variability in RNA expression (interactions among *trans* and *cis* factors and environment). Transcriptome data from mature leaf tissues yielded higher prediction accuracy for most of the phenotypes as compared to the young developing tissues from shoots, roots, or kernels tissue, highlighting the variability in expression regulation across different tissues [26]. Therefore, gene expression in different tissues may not capture the same functional information. Most of the phenotypes in this study were measured under field conditions in mature tissues or kernels in different seasons (e.g., flowering traits, agronomic/field traits). Therefore, mature leaf tissue’s expression measured in the field should be “closer” to these phenotypes, allowing higher prediction accuracy, compared to expression at the seedling stage measured under controlled conditions. With HARE, the contextual issue was less pronounced resulting in more stable prediction accuracy from any of these tissues.

### Baseline for comparison of genomic prediction models is important

To determine if haplotype expression carries functional information in addition to haplotype structure in genomic prediction, we used genetic signals produced by random HARE as a baseline for our genomic prediction models. Prior studies have used different baselines to assess genomic prediction models. For example, Westwhues et al. [42] used traditional pedigree BLUP as a baseline for predicting seven complex traits in hybrids as compared to using genomic sequence, metabolomes, transcriptome or a combinations of these; Azodi et al. [2] used the first five principal components in the marker data and compared these with the genomic and transcriptome data to predict three traits; and Li et al. [5] used a genomic BLUP model with SNP data and compared it to extensions with additional endophenotypes in the model to predict nine traits. In this study, randomly permuting the HARE estimates 100 times while preserving haplotype structure, allowed us to assess the accuracy of genomic prediction in the absence of functional information in haplotype values, and to directly test the significance of HARE over haplotype structure. The significantly higher prediction accuracy of HARE affirmed that HARE carried functional information beyond haplotype structure.

### HARE captured functional information beyond haplotype structure

The benefits of using haplotypes over SNPs, and transcriptomes over SNPs in genomic prediction has been highlighted in earlier studies [4,5,7–10]. Our study here integrated both haplotypes and transcriptome information (as HARE) in the prediction of complex traits. Haplotypes can capture the interaction and epistasis occurring in genic regions, which cannot be captured by SNP data alone [4–6,42]. Another issue in genomic prediction models is overparameterization, where there are more predictors than observations [2,43]. By using transcriptome data rather than SNPs as predictors, the feature dimension can be reduced from millions to thousands, which should make the model more transferable by addressing the curse of dimensionality. This approach could be further improved by functional annotations about gene expression, which could give more or less importance to some genes in HARE, based on prior knowledge about their effect on phenotypes.

### Tapping into a new source of functional information using HARE

Here, we presented a novel method for imputing HARE using the Practical Haplotype Graph (PHG) and a mixed model approach (Fig 1, 8). Measuring the transcriptome in multiple tissues for every population is expensive, while imputing expression is easier and cost effective. Here we used existing transcriptomes profiled in 7 diverse tissues of the Goodman Association panel consisting of 280 diverse lines to get HARE estimates in NAM founders’ haplotypes. HARE estimates were then imputed in the NAM panel consisting of 5000 lines using the PHG. Imputing expression was not only cheaper, it also contributed to more robust genomic predictions. Other methods for predicting expression from genomic sequence were previously reported however these methods are less accurate [15,44]. Compared to these methods, our approach here requires sparse sequencing data only to obtain haplotypes by the PHG and expression in some genotypes. Therefore, it was less computation intensive and more cost-effective than approaches based on deep neural networks applied to complete sequencing data.

**Fig 8.**
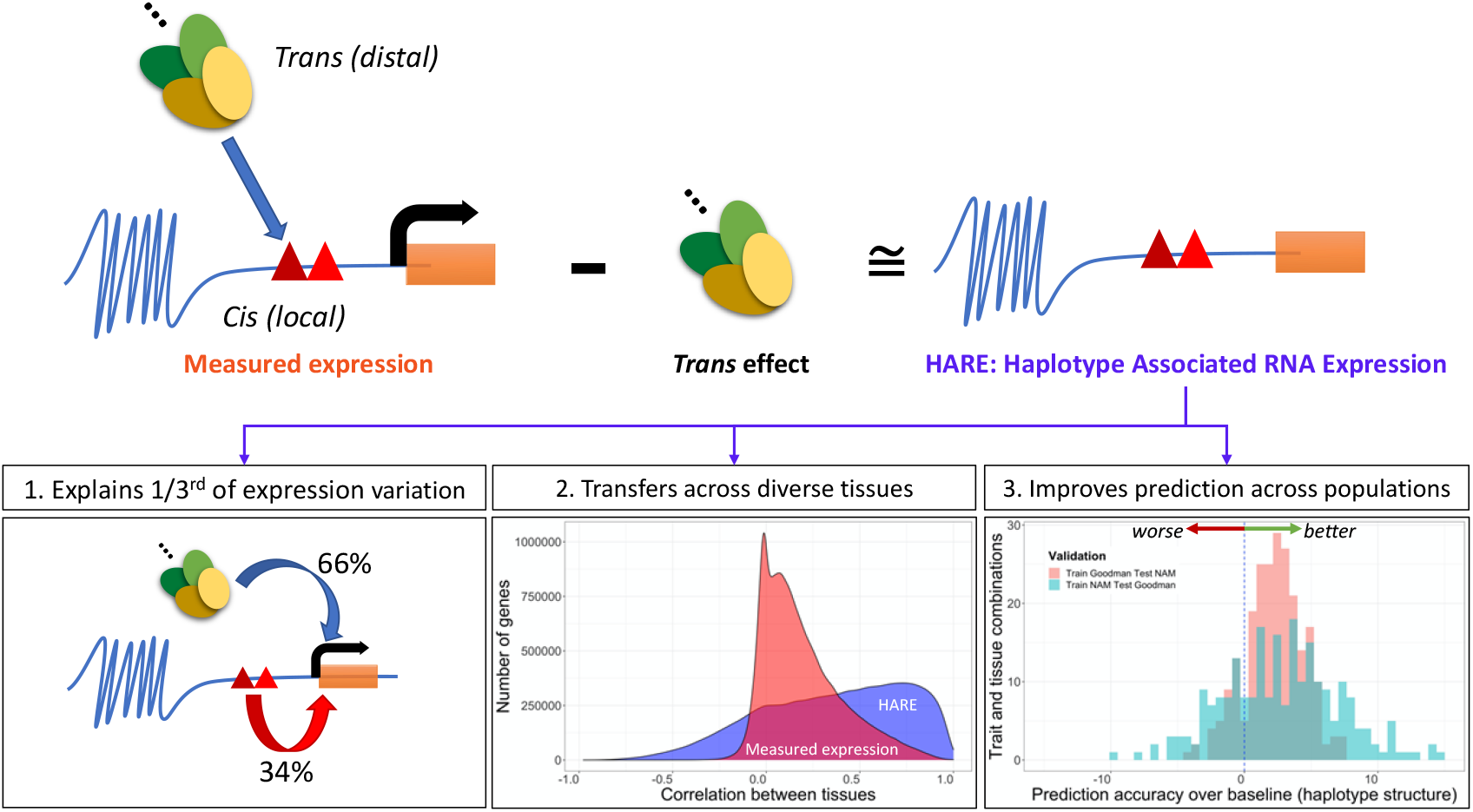
Graphical Summary of the Study. *Cis* haplotype associated RNA expression (HARE) was obtained from subtracting *trans* effects from measured expression. 1. The *cis* haplotypes explained one-third of the variation in expression. 2. The HARE estimates were highly transferrable across tissues compared to measured expression. 3. HARE improved prediction within and across populations in maize.

### Conclusion

We showed that by leveraging the diverse high-quality assemblies through a haplotype graph, we can impute *cis* Haplotype Associated RNA Expression in diverse panels. By showing higher transferability across tissues and moderate correlation with measured expression, we have demonstrated that imputing HARE could generate more stable measurements of gene expression across biological contexts. Also, we have demonstrated that HARE estimates could improve genomic prediction for most complex traits in maize over haplotype structure or measured expression. The important consideration in many genomic prediction and transcriptome studies is the cost of generating the genomics and transcriptomics data. Our approach here utilizes sparse sequencing data to obtain haplotypes and impute expression on those haplotypes using the expression measured in some related genotypes.

Future experiments could aim at using HARE to predict across species, using gene regulatory networks to provide priors for key genes and trait combinations, refining expression estimates by adding machine learning models that predict expression from DNA sequence, and finally more explicitly modeling tissue by genotype interactions. However, because a substantial proportion of expression is stable across tissues and it can be estimated by variance partitioning, HARE can be applied to successfully improve phenotypic prediction at modest cost.

## Materials and methods

### Phenotypic data

Two maize panels were evaluated for prediction accuracy: the US Nested Association Mapping (NAM) panel and the Goodman Association panel representing the genetic diversity among maize elite inbred lines. The NAM panel was developed from 25 parents crossed to a common parent B73 and selfed to obtain 200 homozygous recombinant inbred lines (RILs) from each cross, as described in McMullen et al. [20] and Gage et al. [21]. The Goodman Association panel represents the global diversity of inbred lines in public maize breeding programs, including ~282 genotypes from tropical and temperate regions, sweet corn, and popcorn lines [19]. The 25 NAM founders are part of the Goodman Association panel, so we excluded them from the Goodman Association panel set for cross-panel prediction.

We evaluated genomic prediction models for 26 traits belonging to different groups: flowering, morphology, yield-related, kernel composition, and disease (Table 1). These traits were chosen from three publications where they were jointly phenotyped in the two panels [22–25]. Phenotypic evaluations for these traits were performed in 2006 and 2007 across 11 environments, though not all traits were measured in all environments. The field experiments were conducted using an incomplete block alpha lattice design. The phenotypic values were best linear unbiased predictors (BLUPs), details on the phenotypic measurement and BLUP calculation are presented in the respective studies (Table 1).

### PHG database for NAM and Goodman Association panels

Details on the Practical Haplotype Graph (PHG) were described in [27]. In brief, the database consisted of the genomes of 26 NAM parents and one additional stiff stalk inbred B104. The genomes were divided into reference ranges, where the edges of each reference range were defined by gene boundaries in B73 RefGen_v5. A total of 71,354 reference ranges were identified, where half of them were genic regions. The genotyping-by-sequencing (GBS) reads from NAM RILs [45] and the Goodman Association panel [46] were mapped to the PHG database to identify the haplotypes in these populations based on the 27 genomes in the PHG. The SNP calls thus generated were tested for error rate and heterozygosity, imputation accuracy as presented in the original publication [27].

### Haplotype ID analysis

For each line in the NAM and Goodman panels, a haplotype ID was obtained in each reference region from the PHG database using function pathsForMethod in the rPHG package in R (Bradbury et al., in prep). Since the reference ranges included both genic and intergenic regions, the ranges were filtered to yield only the genic reference ranges based on the B73 RefGen_v5 annotations. Shannon entropy of haplotypes counts in the filtered genic reference range were calculated using the R entropy package using the maximum likelihood method [47].

### Gene expression data

Gene expression data were obtained from Kremling et al. [26]. Details on sampling and expression quantification were presented in the original publication. Seven different tissues (germinating seedlings root: GRoot; germinating seedlings shoot: GShoot; two centimeters from base of leaf 3: L3Base; two centimeters from tip of leaf 3: L3Tip; mature mid-leaf tissue sampled during mid-day: LMAD; mature mid-leaf tissue sampled during mid-night: LMAN; and developing kernels harvested after 350 GDD after pollination: Kern) were included in the analysis. Along with the expression from 7 different tissues, maximum expression and average expression per gene were calculated using a custom script in R.

The gene expression data from version 3 were obtained in version 5 of the genome by mapping B73 v3 genes to the v5 reference genome. Genes that did not map or mapped in multiple positions were removed from the analysis. The final genic haplotype matrix included ~18,000 genes with one-to-one correspondence between the two versions of the genome.

### Variance partition in gene expression

The variance components in gene expression were estimated using the R package regress and genetic values were obtained by solving mixed model equations by restricted maximum likelihood (REML) [48]. We fit a linear mixed model for each gene to partition variance into the fraction attributable to the genic reference range (haplotypes representing *cis* effects) with and without controlling for *trans* effects. The effects of haplotypes in the genic reference range were fit as fixed or random as described below. The statistical models for variance partition were:

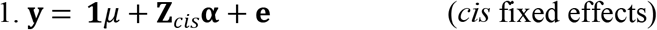

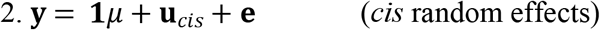

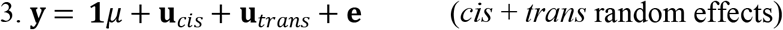

where **y** is the RNA expression at a given gene, **Z**_*cis*_ is the design matrix for the gene’s *cis* haplotypes, **α** is the vector of fixed effects of *cis* haplotypes on gene expression, 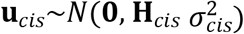 is the vector of *cis* haplotypic effects 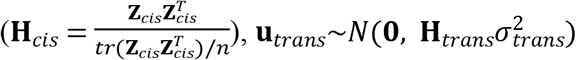 is the vector of *trans* haplotypic effects as captured by the design matrix **Z**_*trans*_ for haplotypes at all genes 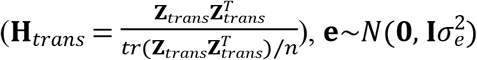 is the vector of errors, *n* is the number of lines in the panel(s), and *tr* is the trace operator (sum of diagonal elements).

The proportion of variance explained by *cis* and *trans* components was estimated from model 3. The *cis* haplotype variance was estimated as 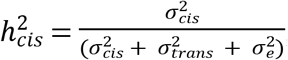, and *trans* variance was estimated as 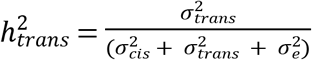. The proportion of heritable variance is the total proportion of variance explained by *cis* and *trans* estimated as 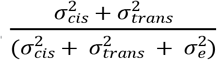, and *cis* portion of heritable variance estimated as 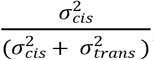.

### Haplotype associated RNA expression (HARE)

The HARE estimates were obtained using the regress package in R and genetic values were obtained by solving mixed model equations by REML [48]. Models 1, 2, and 3 were used to obtain HARE estimates for each haplotype in all genic regions.

Expression matrices were generated for genes in the Goodman panel and NAM based on the 27 haplotypes from NAM parents and B104. Missing haplotype expression was imputed using mean imputation using a custom script in R. The HARE expression matrix was compared with the measured expression matrix in the Goodman panel by pairwise correlation of genes using the *cor* function in R. Pairwise correlation was calculated between measured expression and HARE estimated across all genes. Similarly, transferability across tissues were assessed by pairwise correlation of genes across all 21 different combinations of 7 tissues for both measured expression and HARE.

### Genomic prediction model and model performance

The genomic prediction model was fit using ridge regression using the glmnet package in R [55] and the optimal value of the regularization parameter λ was determined by minimum mean squared error in 10-fold cross-validation.

For a given set of *n* individuals and *p* genes, the following linear model was fit:

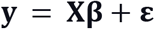

Where **y** is a *n*-vector of phenotypic values, **X** is the *n* x *p* matrix of expression values (measured expression or HARE estimates), ***β*** is the *p*-vector of effects of expression on phenotypes and 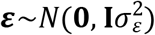 is the vector of errors.

### Assessment of genomic prediction ability

Part of the signal in genomic prediction by HARE may have been due to the sharing of haplotypes in these populations. Therefore, we established a baseline for genomic prediction by using random HARE estimates for haplotypes while preserving the haplotype structure (random HARE). For random HARE, the HARE estimates were permuted at the haplotype level, so that each gene had the same HARE estimates, but HARE estimates were randomly matched to haplotypes. For example, if lines 1 and 2 carried the same haplotype at gene *j*, after permutation, both lines got the same random value for imputed expression (Fig 1). The significance of using HARE over random HARE was assessed by using a Monte Carlo procedure with 100 random permutations of HARE [49]. P-values are calculated as 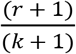 where *k* = 100 is the total number of permutations and r is the number of permutations with accuracy greater than using HARE (accuracy of random HARE greater than accuracy of HARE).

The prediction accuracy of the genomic prediction models was defined as the Pearson correlation coefficient between the observed trait values (**y**) and predicted values (**ŷ**) in each of the test sets using the *cor* function in R.

### Within-panel prediction

In within-panel prediction, the prediction was carried out only in the Goodman panel using the measured expression, HARE, and random HARE (representing haplotype structure). We used a 5-fold cross-validation procedure for all data sets. The panel was randomly partitioned into 80% training set and 20% testing set. Partitions were repeated 20 times. Pearson correlation was individually calculated in each of the 20 partitions and averaged over partitions to test for significance. For a single trait and tissue combination, the model was run for 2000 times for random HARE and 20 times for HARE and measured expression.

### Cross-panel prediction

For cross-population prediction, the model was either trained in NAM and tested in the Goodman panel or vice versa for all traits. The 25 NAM founders are part of the Goodman Association panel, so we excluded them from the Goodman Association panel set for cross-panel prediction. To account for sample size differences (5000 in NAM *versus* 250 in the Goodman panel), 20 random subsets of NAM equivalent to the size of Goodman panel were created, taking 10 RILs from each family. The model was trained in the 20 random subsets of NAM RILs and predicted in the Goodman panel and vice versa for three sample traits: Days to anthesis (DTA), Days to silking (DTS), and Plant height (PH). The prediction accuracy was averaged across 20 random subsets.

## Acknowledgments

This study was conducted using the gene expression data collected in seven tissues of Goodman Association Panel (Kremling et al. [26]) and the Practical Haplotype Graph database for the Goodman Association Panel and Nested Association Mapping Panel (Valdes Franco et al. [27]). We thank Sara Miller for copy editing.

---

S1 Fig. Phenotypic distribution of 26 traits in NAM and the Goodman Association panel

S2 Fig. a. Haplotype counts distribution in the Goodman association panel (left) and NAM panel (right) across all genic reference ranges. b. Haplotype entropy in Goodman (left) and NAM (right) panel in each reference range. Median haplotype counts were 8 and 100 in the Goodman panel and NAM, respectively, resulting in higher entropy in the Goodman panel as compared to NAM. Entropy was calculated from haplotype counts in each reference region.

S3 Fig. Correlation distribution of expression between tissues. The four panels represent HARE estimates from models 1 (*cis* only fixed), 2 (*cis* only random), and 3 (*cis* random while accounting for *trans*), as well as measured expression. The different color lines in each panel represent 21 different combinations of the 7 different tissues as labeled on the right: germinating seedlings root (GRoot), germinating seedlings shoot (GShoot), two cm from base of leaf 3 (L3Base), two cm from tip of leaf 3 (L3Tip), mature mid-leaf tissue sampled during mid-day (LMAD), mature mid-leaf tissue sampled during mid-night (LMAN), and developing kernels harvested after 350 growing degree days after pollination (Kern). The imputed expression from models was highly correlated between tissues when compared to the measured transcript expression. In all panels, closely related tissues like matured mid-leaf tissue expression sampled during mid-day (LMAD) and matured mid-leaf tissue expression sampled during mid-night (LMAD) were highly correlated.

S4 Fig. Correlation distribution of *trans* components of expression between tissues. The different color lines in each panel represent 21 different combinations of the 7 different tissues as labeled on the right: germinating seedlings root (GRoot), germinating seedlings shoot (GShoot), two cm from base of leaf 3 (L3Base), two cm from tip of leaf 3 (L3Tip), mature mid-leaf tissue sampled during mid-day (LMAD), mature mid-leaf tissue sampled during mid-night (LMAN), and developing kernels harvested after 350 growing degree days after pollination (Kern). Similar to measured transcript expression, closely related tissues like matured leaf expression during the day (LMAD) and matured leaf expression during the night (LMAD) were highly correlated.

S5 Fig. Haplotype associated RNA expression (HARE) was highly correlated across tissues as compared to measured transcript expression. Different colors represent HARE imputed from three statistical models: Model 1 (*cis* fixed effect), 2 (*cis* random effect), and 3 (*cis* + *trans* random effects), and measured transcript expression. The distribution is the pairwise correlation of ~8000 highly expressed genes across 21 different combinations from 7 different tissues.

S6 Fig. Prediction accuracy using HARE from model 1, 2, and 3 (see methods) for predicting three different traits: Days to Anthesis (DTA), Days to Silking (DTS), and Plant Height (PH) using a) model trained in NAM and tested in Goodman b) model trained in Goodman and tested in NAM. The different symbols represent HARE from different tissues: germinating seedlings shoot (GShoot), developing kernels harvested after 350 growing degree days after pollination (Kern), 2 cm from base of leaf 3 (L3Base), and mature mid-leaf tissue sampled during mid-day (LMAD).

S7 Fig. Within-panel prediction accuracy in the Goodman panel using HARE (red dot), 100 random HARE (box plot), and measured expression (blue dot) from individual tissues or all tissues integrated as mean or maximum expression. Individual tissues included: germinating seedlings root (GRoot), germinating seedlings shoot (GShoot), two cm from base of leaf 3 (L3Base), two cm from tip of leaf 3 (L3Tip), mature mid-leaf tissue sampled during mid-day (LMAD), mature mid-leaf tissue sampled during mid-night (LMAN), and developing kernels harvested after 350 growing degree days after pollination (Kern). The model was trained in 80% of the panel and tested in the remaining 20%.

S8 Fig. Change in prediction accuracy using HARE over the mean of random expression (blue dashed line) from five different tissues: germinating seedlings root (GRoot), two cm from base of leaf 3 (L3Base), mature mid-leaf tissue sampled during mid-day (LMAD), mature mid-leaf tissue sampled during mid-night (LMAN), and developing kernels harvested after 350 growing degree days after pollination (Kern). Genomic prediction models were (a) trained in the Goodman panel and tested in 20 subsets of NAM, (b) trained in 20 subsets of NAM and tested in the Goodman panel. The subsets of NAM were generated by randomly selecting 10 genotypes from each family resulting in a total of 250 genotypes (see methods). Accuracy was averaged over the 20 random subsets before determining significance. The black shapes represent statistically significant differences at P-values <0.05 and red shapes represent no statistical significance. P-values were calculated using a Monte Carlo procedure.

S1 Table. Prediction accuracy of 26 complex traits in the Goodman Association panel using HARE from seven diverse tissues: germinating seedlings root (GRoot), germinating seedlings shoot (GShoot), 2 cm from base of leaf 3 (L3Base), two cm from tip of leaf 3 (L3Tip), mature mid-leaf tissue sampled during mid-day (LMAD), mature mid-leaf tissue sampled during mid-night (LMAN), and developing kernels harvested after 350 growing degree days after pollination (Kern), mean, and maximum expression of genes across all tissues. P value (high) and P value (low) were calculated using a Monte Carlo procedure to test if the accuracy using HARE was significantly higher or lower than random HARE. Models were trained in NAM and tested in Goodman Association panel.

S2 Table. Prediction accuracy of 26 complex traits in NAM using HARE from 7 diverse tissues: germinating seedlings root (GRoot), germinating seedlings shoot (GShoot), 2 cm from base of leaf 3 (L3Base), 2 cm from tip of leaf 3 (L3Tip), mature mid-leaf tissue sampled during mid-day (LMAD), mature mid-leaf tissue sampled during mid-night (LMAN), and developing kernels harvested after 350 growing degree days after pollination (Kern), mean, and maximum expression of genes across all tissues. P value (high) and P value (low) were calculated using a Monte Carlo procedure to test if the accuracy using HARE was significantly higher or lower than random HARE. Models were trained in Goodman Association panel and tested in NAM.

## References

1. Meuwissen THE, Hayes BJ, Goddard ME. Prediction of total genetic value using genome-wide dense marker maps. Genetics. 2001;157: 1819–1829.

2. Azodi CB, Pardo J, VanBuren R, de los Campos G, Shiu S-H. Transcriptome-Based Prediction of Complex Traits in Maize. Plant Cell. 2020;32: 139–151. doi:10.1105/tpc.19.00332

3. Xu Y, Xu C, Xu S. Prediction and association mapping of agronomic traits in maize using multiple omic data. Heredity. 2017;119: 174–184. doi:10.1038/hdy.2017.27

4. Guo Z, Magwire MM, Basten CJ, Xu Z, Wang D. Evaluation of the utility of gene expression and metabolic information for genomic prediction in maize. Theor Appl Genet. 2016;129: 2413–2427. doi:10.1007/s00122-016-2780-5

5. Li Z, Gao N, Martini JWR, Simianer H. Integrating Gene Expression Data Into Genomic Prediction. Front Genet. 2019;10. doi:10.3389/fgene.2019.00126

6. Schrag TA, Westhues M, Schipprack W, Seifert F, Thiemann A, Scholten S, et al. Beyond Genomic Prediction: Combining Different Types of *omics* Data Can Improve Prediction of Hybrid Performance in Maize. Genetics. 2018;208: 1373–1385. doi:10.1534/genetics.117.300374

7. Hayes BJ, Chamberlain AJ, McPARTLAN H, Macleod I, Sethuraman L, Goddard ME. Accuracy of marker-assisted selection with single markers and marker haplotypes in cattle. Genet Res. 2007;89: 215–220. doi:10.1017/S0016672307008865

8. Hess M, Druet T, Hess A, Garrick D. Fixed-length haplotypes can improve genomic prediction accuracy in an admixed dairy cattle population. Genet Sel Evol. 2017;49: 54. doi:10.1186/s12711-017-0329-y

9. Won S, Park J-E, Son J-H, Lee S-H, Park BH, Park M, et al. Genomic Prediction Accuracy Using Haplotypes Defined by Size and Hierarchical Clustering Based on Linkage Disequilibrium. Front Genet. 2020;11. doi:10.3389/fgene.2020.00134

10. Schopp P, Müller D, Technow F, Melchinger AE. Accuracy of Genomic Prediction in Synthetic Populations Depending on the Number of Parents, Relatedness, and Ancestral Linkage Disequilibrium. Genetics. 2017;205: 441–454. doi:10.1534/genetics.116.193243

11. Signor SA, Nuzhdin SV. The Evolution of Gene Expression in cis and trans. Trends Genet. 2018;34: 532–544. doi:10.1016/j.tig.2018.03.007

12. Albert FW, Bloom JS, Siegel J, Day L, Kruglyak L. Genetics of trans-regulatory variation in gene expression. eLife. 2018;7: 7:e3547. doi:10.7554/eLife.35471

13. Tu X, Mejía-Guerra MK, Valdes Franco JA, Tzeng D, Chu P-Y, Shen W, et al. Reconstructing the maize leaf regulatory network using ChIP-seq data of 104 transcription factors. Nat Commun. 2020;11: 5089. doi:10.1038/s41467-020-18832-8

14. Wittkopp PJ, Haerum BK, Clark AG. Evolutionary changes in cis and trans gene regulation. Nature. 2004;430: 85–88. doi:10.1038/nature02698

15. Washburn JD, Mejia-Guerra MK, Ramstein G, Kremling KA, Valluru R, Buckler ES, et al. Evolutionarily informed deep learning methods for predicting relative transcript abundance from DNA sequence. Proc Natl Acad Sci. 2019;116: 5542–5549. doi:10.1073/pnas.1814551116

16. Grundberg E, Adoue V, Kwan T, Ge B, Duan QL, Lam KCL, et al. Global Analysis of the Impact of Environmental Perturbation on cis-Regulation of Gene Expression. Gibson G, editor. PLoS Genet. 2011;7: e1001279. doi:10.1371/journal.pgen.1001279

17. Schnable PS, Ware D, Fulton RS, Stein JC, Wei F, Pasternak S, et al. The B73 maize genome: complexity, diversity, and dynamics. Science. 2009;326: 1112–1115. doi:10.1126/science.1178534

18. Wallace JG, Bradbury PJ, Zhang N, Gibon Y, Stitt M, Buckler ES. Association Mapping across Numerous Traits Reveals Patterns of Functional Variation in Maize. PLOS Genet. 2014;10: e1004845. doi:10.1371/journal.pgen.1004845

19. Flint-Garcia SA, Thuillet A-C, Yu J, Pressoir G, Romero SM, Mitchell SE, et al. Maize association population: a high-resolution platform for quantitative trait locus dissection: High-resolution maize association population. Plant J. 2005;44: 1054–1064. doi:10.1111/j.1365-313X.2005.02591.x

20. McMullen MD, Kresovich S, Villeda HS, Bradbury P, Li H, Sun Q, et al. Genetic Properties of the Maize Nested Association Mapping Population. Science. 2009;325: 737–740. doi:10.1126/science.1174320

21. Gage JL, Monier B, Giri A, Buckler ES. Ten Years of the Maize Nested Association Mapping Population: Impact, Limitations, and Future Directions. Plant Cell. 2020;32: 2083–2093. doi:10.1105/tpc.19.00951

22. Hung H-Y, Shannon LM, Tian F, Bradbury PJ, Chen C, Flint-Garcia SA, et al. ZmCCT and the genetic basis of day-length adaptation underlying the postdomestication spread of maize. Proc Natl Acad Sci. 2012;109: E1913–E1921. doi:10.1073/pnas.1203189109

23. Kump KL, Bradbury PJ, Wisser RJ, Buckler ES, Belcher AR, Oropeza-Rosas MA, et al. Genome-wide association study of quantitative resistance to southern leaf blight in the maize nested association mapping population. Nat Genet. 2011;43: 163–168. doi: 10.1038/ng.747

24. Cook JP, McMullen MD, Holland JB, Tian F, Bradbury P, Ross-Ibarra J, et al. Genetic Architecture of Maize Kernel Composition in the Nested Association Mapping and Inbred Association Panels. PLANT Physiol. 2012;158: 824–834. doi:10.1104/pp.111.185033

25. Peiffer JA, Romay MC, Gore MA, Flint-Garcia SA, Zhang Z, Millard MJ, et al. The Genetic Architecture Of Maize Height. Genetics. 2014;196: 1337–1356. doi:10.1534/genetics.113.159152

26. Kremling KAG, Chen S-Y, Su M-H, Lepak NK, Romay MC, Swarts KL, et al. Dysregulation of expression correlates with rare-allele burden and fitness loss in maize. Nature. 2018;555: 520–523. doi:10.1038/nature25966

27. Franco JAV, Gage JL, Bradbury PJ, Johnson LC, Miller ZR, Buckler ES, et al. A Maize Practical Haplotype Graph Leverages Diverse NAM Assemblies. Genomics; 2020 Aug. doi:10.1101/2020.08.31.268425

28. Liu X, Li YI, Pritchard JK. Trans Effects on Gene Expression Can Drive Omnigenic Inheritance. Cell. 2019;177: 1022–1034.e6. doi:10.1016/j.cell.2019.04.014

29. GTEx Consortium. Genetic effects on gene expression across human tissues. Nature. 2017;550: 204–213. doi:10.1038/nature24277

30. Lemmon ZH, Bukowski R, Sun Q, Doebley JF. The Role of cis Regulatory Evolution in Maize Domestication. Fraser H, editor. PLoS Genet. 2014;10: e1004745. doi:10.1371/journal.pgen.1004745

31. Lopez-Arboleda WA, Reinert S, Nordborg M, Korte A. Global genetic heterogeneity in adaptive traits. Evolutionary Biology; 2021 Feb. doi:10.1101/2021.02.26.433043

32. Gibson G. Rare and common variants: twenty arguments. Nat Rev Genet. 2012;13: 135–145. doi:10.1038/nrg3118

33. Schmid M, Davison TS, Henz SR, Pape UJ, Demar M, Vingron M, et al. A gene expression map of Arabidopsis thaliana development. Nat Genet. 2005;37: 501–506. doi:10.1038/ng1543

34. Sekhon RS, Lin H, Childs KL, Hansey CN, Buell CR, Leon N de, et al. Genome-wide atlas of transcription during maize development. Plant J. 2011;66: 553–563. doi:10.1111/j.1365-313X.2011.04527.x

35. Melé M, Ferreira PG, Reverter F, DeLuca DS, Monlong J, Sammeth M, et al. The human transcriptome across tissues and individuals. Science. 2015;348: 660–665. doi:10.1126/science.aaa0355

36. Mogil LS, Andaleon A, Badalamenti A, Dickinson SP, Guo X, Rotter JI, et al. Genetic architecture of gene expression traits across diverse populations. Epstein MP, editor. PLOS Genet. 2018;14: e1007586. doi:10.1371/journal.pgen.1007586

37. Stern DL. Perspective: Evolutionary Developmental Biology and the Problem of Variation. Evolution. 2000;54: 1079–1091. doi:https://doi.org/10.1111/j.0014-3820.2000.tb00544.x

38. Wray GA. The evolutionary significance of cis-regulatory mutations. Nat Rev Genet. 2007;8: 206–216. doi:10.1038/nrg2063

39. Initial sequence of the chimpanzee genome and comparison with the human genome. Nature. 2005;437: 69–87. doi:10.1038/nature04072

40. Missra A, Ernest B, Lohoff T, Jia Q, Satterlee J, Ke K, et al. The Circadian Clock Modulates Global Daily Cycles of mRNA Ribosome Loading. Plant Cell. 2015;27: 2582–2599. doi:10.1105/tpc.15.00546

41. Fiévet JB, Dillmann C, de Vienne D. Systemic properties of metabolic networks lead to an epistasis-based model for heterosis. Theor Appl Genet. 2009;120: 463. doi:10.1007/s00122-009-1203-2

42. Westhues M, Schrag TA, Heuer C, Thaller G, Utz HF, Schipprack W, et al. Omics-based hybrid prediction in maize. Theor Appl Genet. 2017;130: 1927–1939. doi:10.1007/s00122-017-2934-0

43. Ramstein GP, Jensen SE, Buckler ES. Breaking the curse of dimensionality to identify causal variants in Breeding 4. Theor Appl Genet. 2019;132: 559–567. doi:10.1007/s00122-018-3267-3

44. Zhou J, Theesfeld CL, Yao K, Chen KM, Wong AK, Troyanskaya OG. Deep learning sequence-based ab initio prediction of variant effects on expression and disease risk. Nat Genet. 2018;50: 1171–1179. doi:10.1038/s41588-018-0160-6

45. Rodgers-Melnick E, Bradbury PJ, Elshire RJ, Glaubitz JC, Acharya CB, Mitchell SE, et al. Recombination in diverse maize is stable, predictable, and associated with genetic load. Proc Natl Acad Sci. 2015;112: 3823–3828. doi:10.1073/pnas.1413864112

46. Romay MC, Millard MJ, Glaubitz JC, Peiffer JA, Swarts KL, Casstevens TM, et al. Comprehensive genotyping of the USA national maize inbred seed bank. Genome Biol. 2013;14: R55. doi:10.1186/gb-2013-14-6-r55

47. Hausser J, Strimmer K, Strimmer M. Package “Entropy.” 2012.

48. Clifford D, McCullagh P. The regress function. R news. 6th ed.: 6–10.

49. North BV, Curtis D, Sham PC. A Note on the Calculation of Empirical P Values from Monte Carlo Procedures. Am J Hum Genet. 2002;71: 439–441.

