## supplemental files for "Haplotype Associated RNA Expression (HARE) Improves Prediction of Complex Traits in Maize"

### Supporting Information:

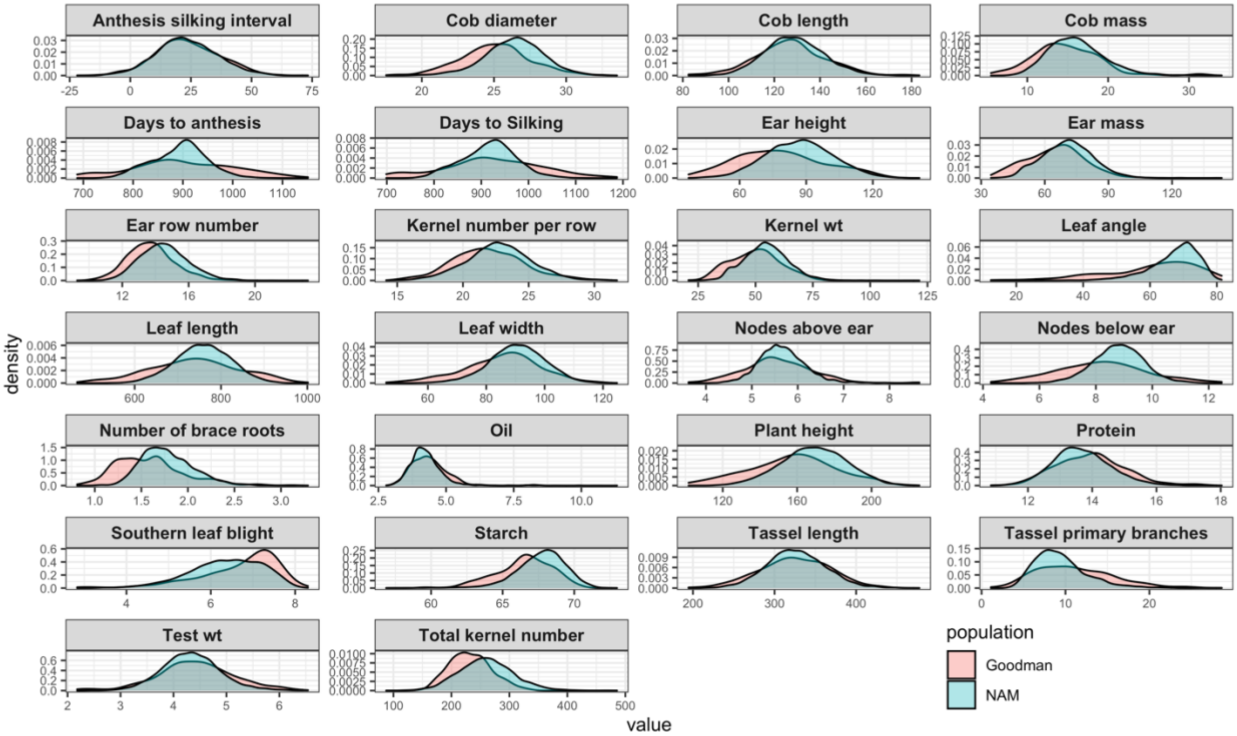


**S1 Fig. Phenotypic distribution of 26 traits in NAM and the Goodman Association panel**

**a.**

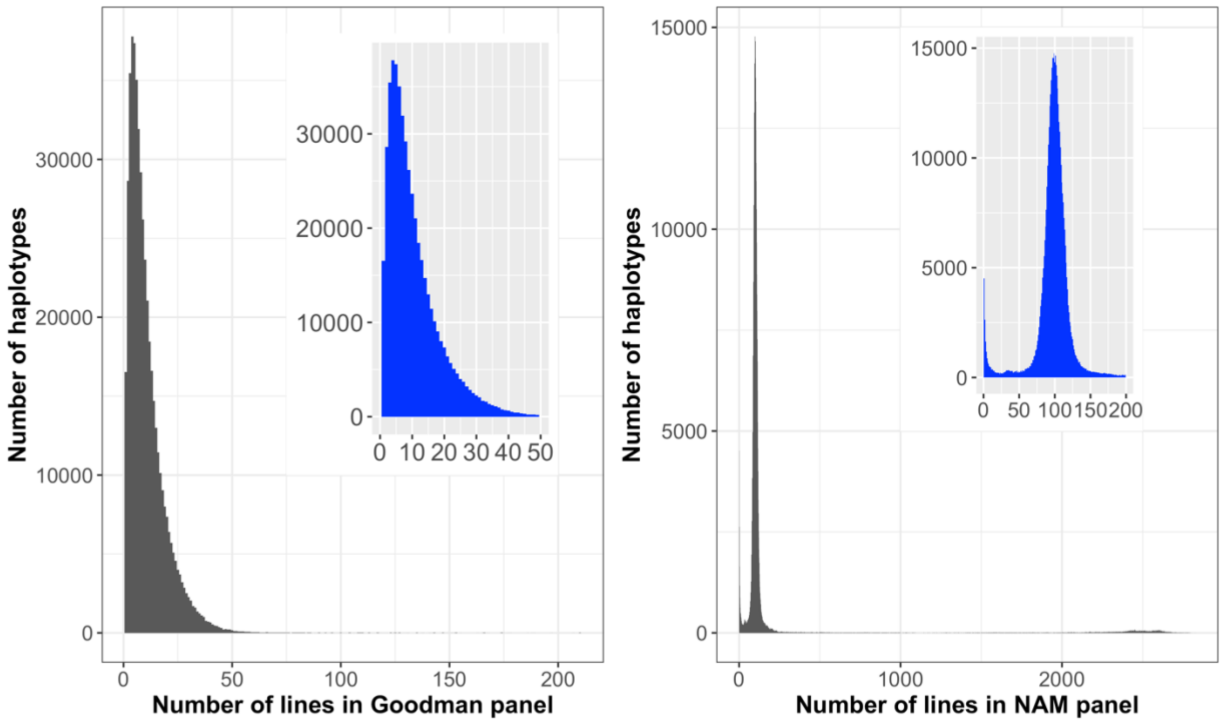

**b.**

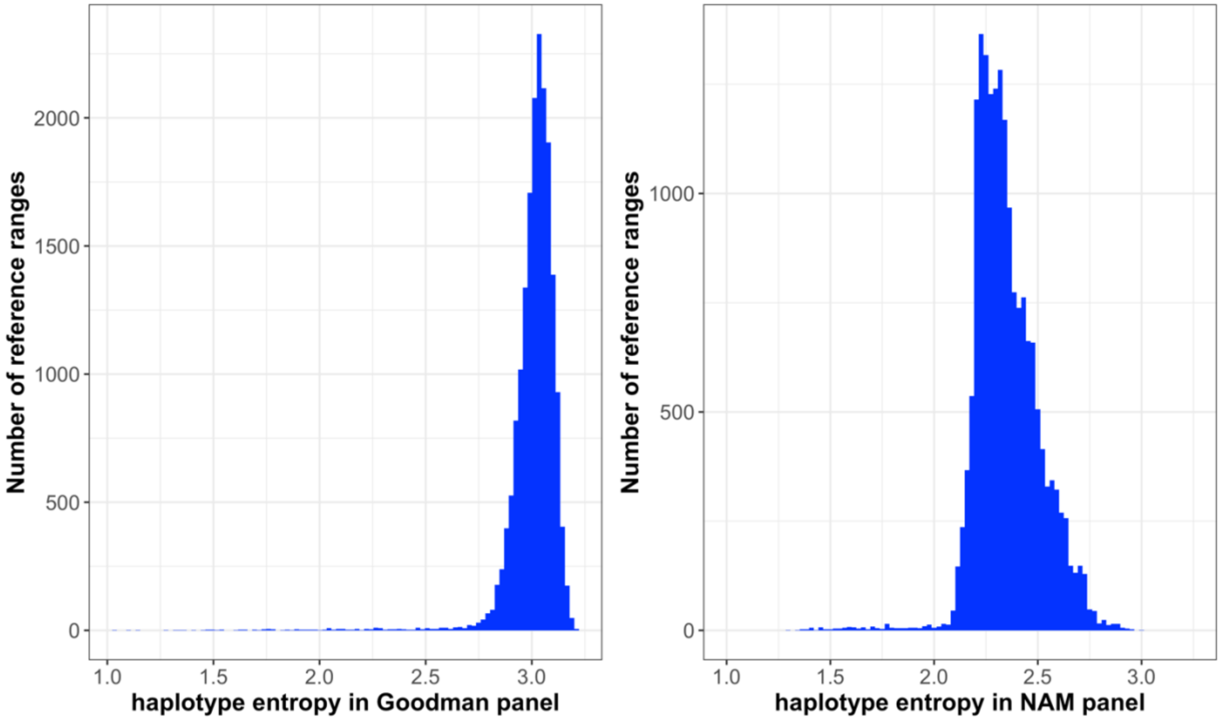


**S2 Fig. a. Haplotype counts distribution in the Goodman association panel (left) and NAM panel (right) across all genic reference ranges. b. Haplotype entropy in Goodman (left) and NAM (right) panel in each reference range. Median haplotype counts were 8 and 100 in the Goodman panel and NAM, respectively, resulting in higher entropy in the Goodman panel as compared to NAM. Entropy was calculated from haplotype counts in each reference region.**

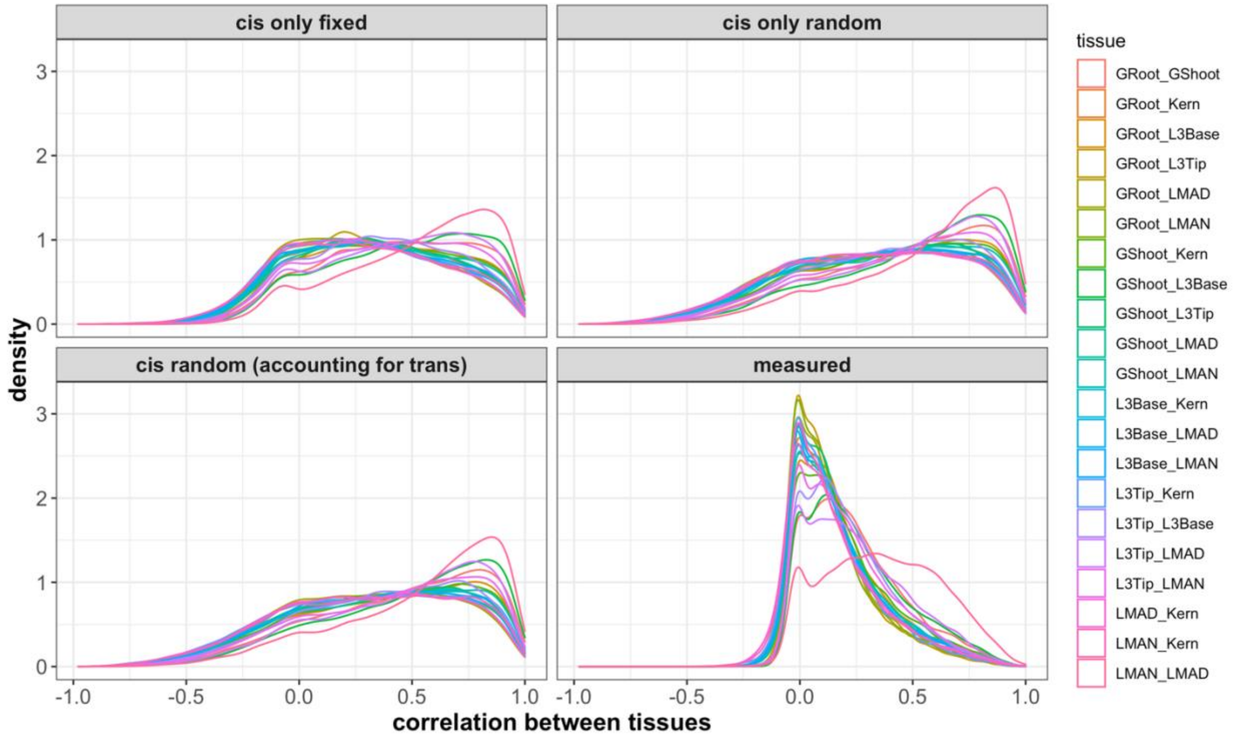


**S3 Fig. Correlation distribution of expression between tissues. The four panels represent HARE estimates from models 1 (*cis* only fixed), 2 (*cis* only random), and 3 (*cis* random while accounting for *trans*), as well as measured expression. The different color lines in each panel represent 21 different combinations of the 7 different tissues as labeled on the right: germinating seedlings root (GRoot), germinating seedlings shoot (GShoot), two cm from base of leaf 3 (L3Base), two cm from tip of leaf 3 (L3Tip), mature mid-leaf tissue sampled during mid-day (LMAD), mature mid-leaf tissue sampled during mid-night (LMAN), and developing kernels harvested after 350 growing degree days after pollination (Kern). The imputed expression from models was highly correlated between tissues when compared to the measured transcript expression. In all panels, closely related tissues like matured mid-leaf tissue expression sampled during mid-day (LMAD) and matured mid-leaf tissue expression sampled during mid-night (LMAD) were highly correlated.**

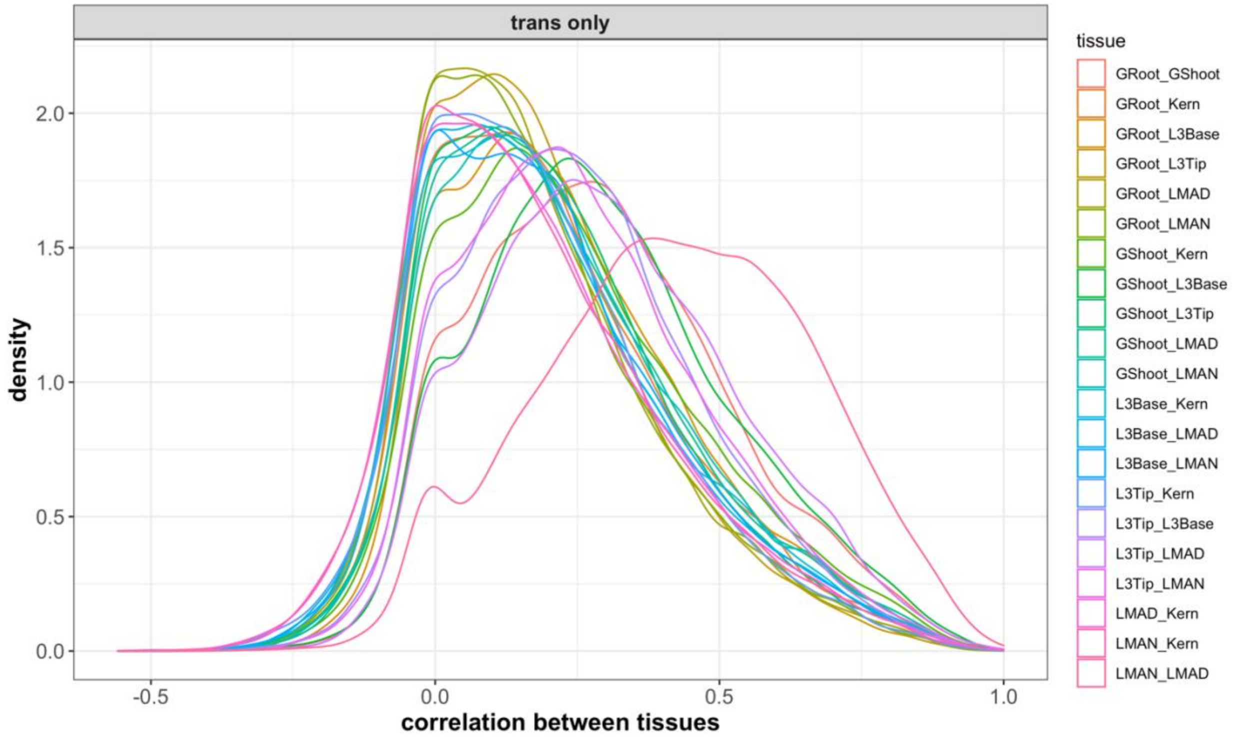


**S4 Fig. Correlation distribution of *trans* components of expression between tissues. The different color lines in each panel represent 21 different combinations of the 7 different tissues as labeled on the right: germinating seedlings root (GRoot), germinating seedlings shoot (GShoot), two cm from base of leaf 3 (L3Base), two cm from tip of leaf 3 (L3Tip), mature mid-leaf tissue sampled during mid-day (LMAD), mature mid-leaf tissue sampled during mid-night (LMAN), and developing kernels harvested after 350 growing degree days after pollination (Kern). Similar to measured transcript expression, closely related tissues like matured leaf expression during the day (LMAD) and matured leaf expression during the night (LMAD) were highly correlated.**

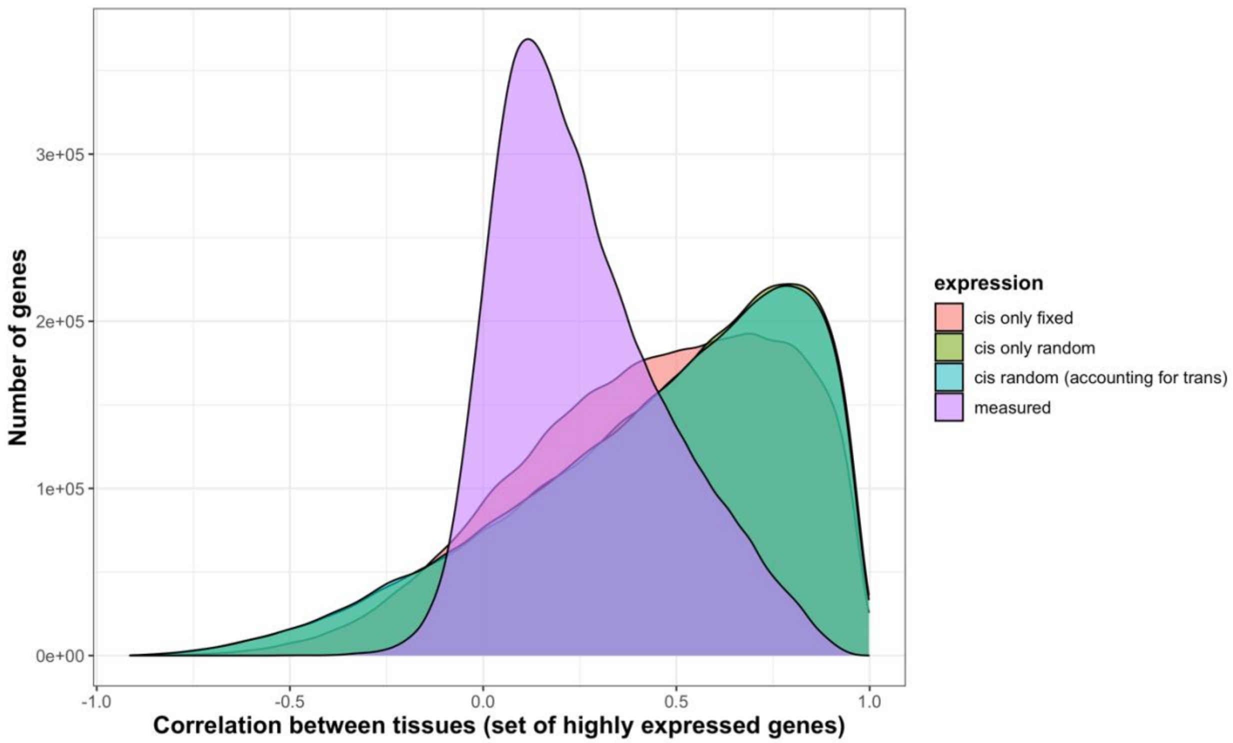


**S5 Fig. Haplotype associated RNA expression (HARE) was highly correlated across tissues as compared to measured transcript expression. Different colors represent HARE imputed from three statistical models: Model 1 (*cis* fixed effect), 2 (*cis* random effect), and 3 (*cis* + *trans* random effects), and measured transcript expression. The distribution is the pairwise correlation of ~8000 highly expressed genes across 21 different combinations from 7 different tissues.**

**a.**

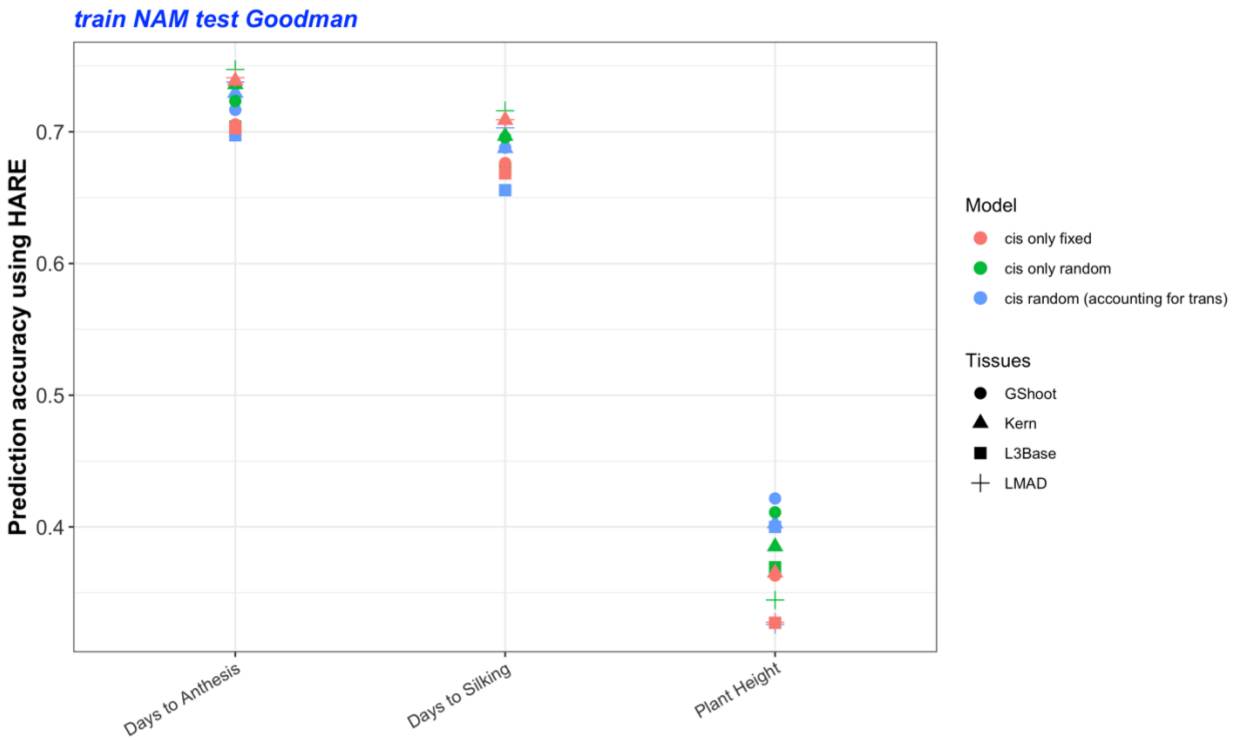


**b.**

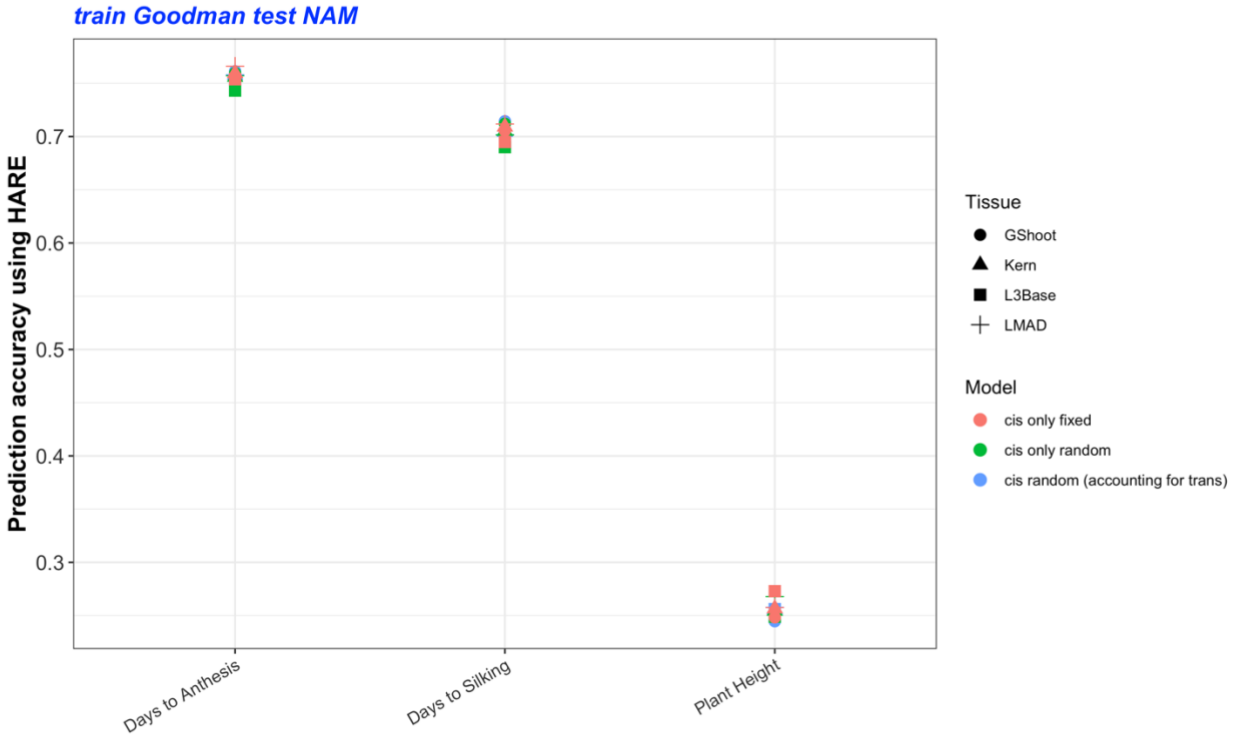


**S6 Fig. Prediction accuracy using HARE from model 1, 2, and 3 (see methods) for predicting three different traits: Days to Anthesis (DTA), Days to Silking (DTS), and Plant Height (PH) using a) model trained in NAM**

**and tested in Goodman b) model trained in Goodman and tested in NAM. The different symbols represent HARE from different tissues: germinating seedlings shoot (GShoot), developing kernels harvested after 350 growing degree days after pollination (Kern), 2 cm from base of leaf 3 (L3Base), and mature mid-leaf tissue sampled during mid-day (LMAD).**

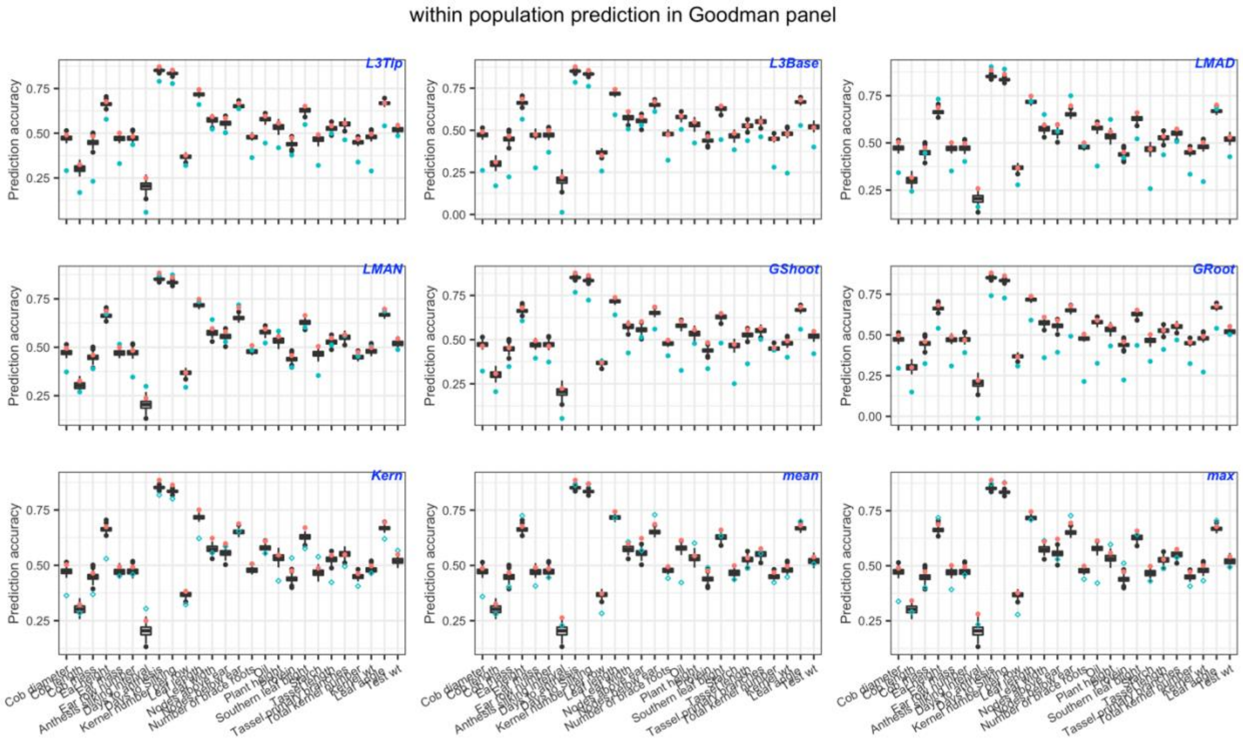


**S7 Fig. Within-panel prediction accuracy in the Goodman panel using HARE (red dot), 100 random HARE (box plot), and measured expression (blue dot) from individual tissues or all tissues integrated as mean or maximum expression. Individual tissues included: germinating seedlings root (GRoot), germinating seedlings shoot (GShoot), two cm from base of leaf 3 (L3Base), two cm from tip of leaf 3 (L3Tip), mature mid-leaf tissue sampled during mid-day (LMAD), mature mid-leaf tissue sampled during mid-night (LMAN), and developing kernels harvested after 350 growing degree days after pollination (Kern). The model was trained in 80% of the panel and tested in the remaining 20%.**

a.

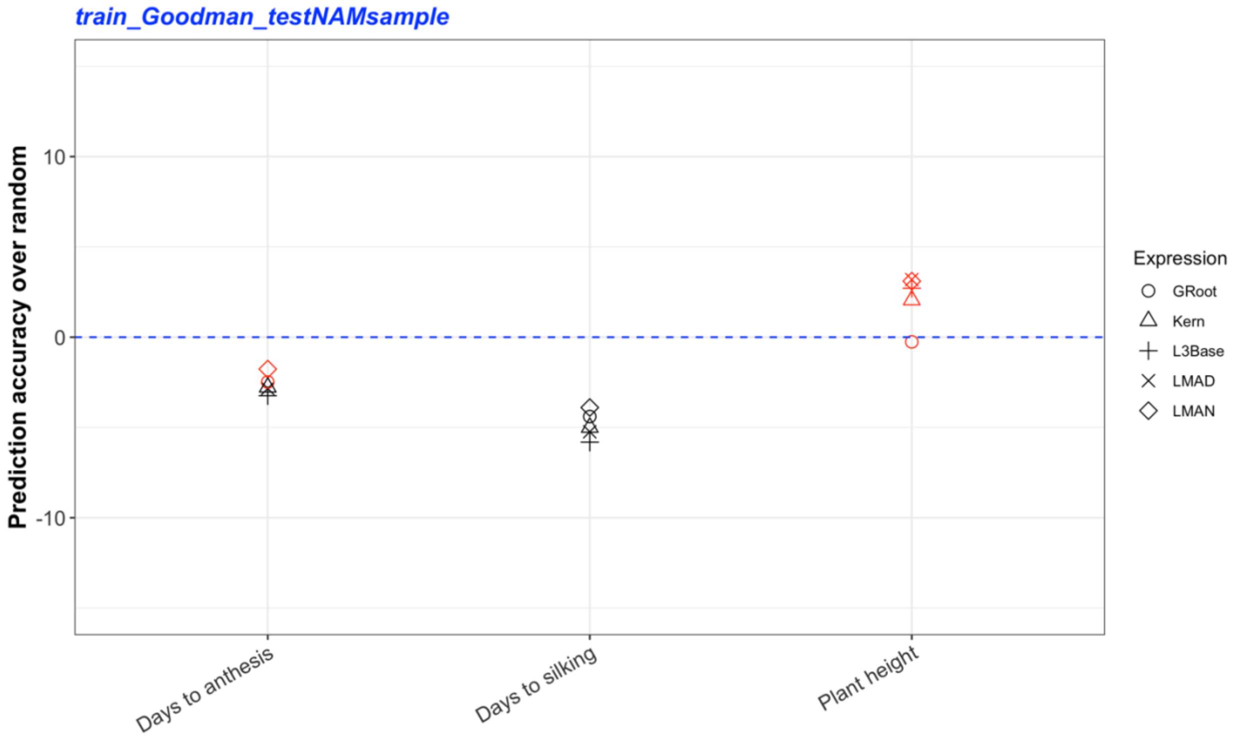


b.

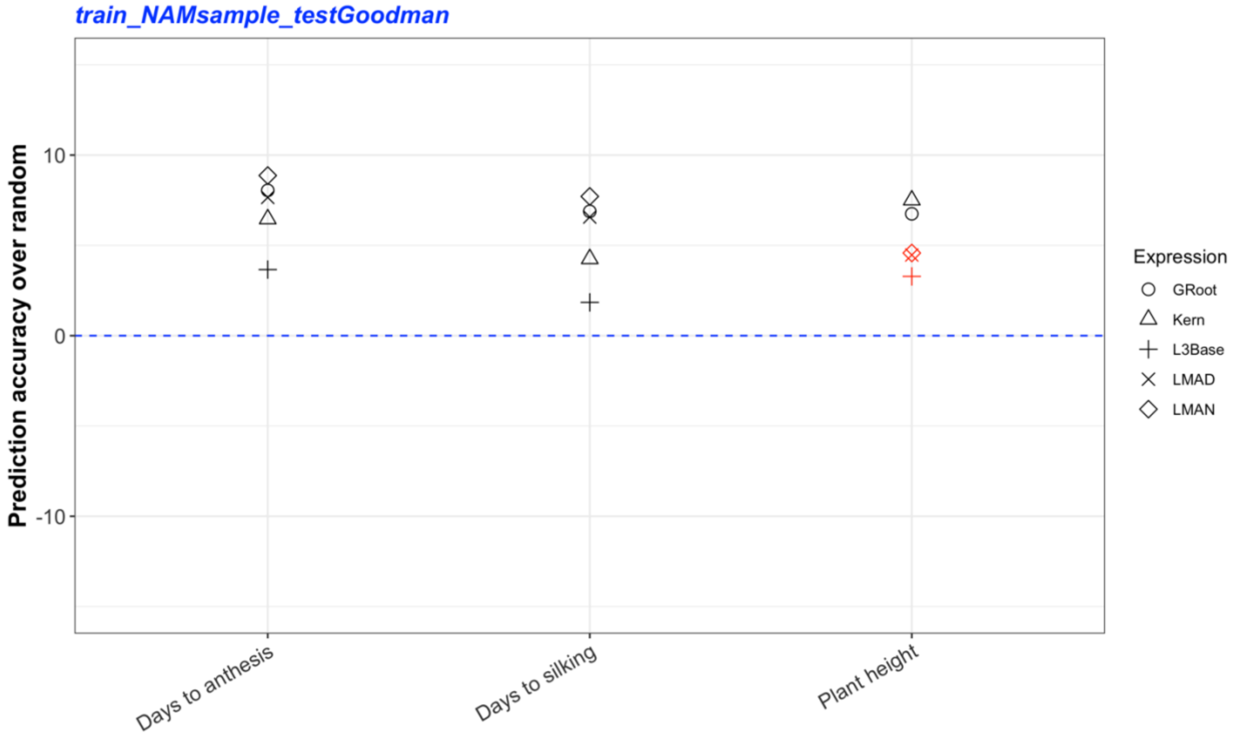


**S8 Fig. Change in prediction accuracy using HARE over the mean of random expression (blue dashed line) from five different tissues: germinating seedlings root (GRoot), two cm from base of leaf 3 (L3Base), mature mid-leaf tissue sampled during mid-day (LMAD), mature mid-leaf tissue sampled during mid-night (LMAN),**

**and developing kernels harvested after 350 growing degree days after pollination (Kern). Genomic prediction models were (a) trained in the Goodman panel and tested in 20 subsets of NAM, (b) trained in 20 subsets of NAM and tested in the Goodman panel. The subsets of NAM were generated by randomly selecting 10 genotypes from each family resulting in a total of 250 genotypes (see methods). Accuracy was averaged over the 20 random subsets before determining significance. The black shapes represent statistically significant differences at P-values <0.05 and red shapes represent no statistical significance. P-values were calculated using a Monte Carlo procedure.**

| Trait | Accuracy using HARE | Mean accuracy using random HARE | P value (high) | P value (low) | Accuracy of HARE over random HARE | Tissues |
|---|---|---|---|---|---|---|
| **Cob diameter** | 0.411 | 0.348 | 0.010 | 1.000 | 6.243 | L3Tip |
| **Cob length** | 0.146 | 0.157 | 0.673 | 0.337 | -1.081 | L3Tip |
| **Cob mass** | 0.302 | 0.266 | 0.129 | 0.881 | 3.657 | L3Tip |
| **Ear height** | 0.559 | 0.473 | 0.010 | 1.000 | 8.620 | L3Tip |
| **Ear mass** | 0.387 | 0.365 | 0.099 | 0.911 | 2.242 | L3Tip |
| **Ear row number** | 0.387 | 0.381 | 0.426 | 0.584 | 0.668 | L3Tip |
| **Anthesis silking interval** | 0.001 | 0.059 | 0.901 | 0.109 | -5.816 | L3Tip |
| **Days to anthesis** | 0.729 | 0.675 | 0.010 | 1.000 | 5.466 | L3Tip |
| **Days to Silking** | 0.693 | 0.642 | 0.010 | 1.000 | 5.057 | L3Tip |
| **Kernel number per row** | 0.194 | 0.195 | 0.574 | 0.436 | -0.084 | L3Tip |
| **Leaf length** | 0.535 | 0.454 | 0.020 | 0.990 | 8.062 | L3Tip |
| **Leaf width** | 0.365 | 0.378 | 0.693 | 0.317 | -1.314 | L3Tip |
| **Nodes above ear** | 0.296 | 0.318 | 0.762 | 0.248 | -2.223 | L3Tip |
| **Nodes below ear** | 0.608 | 0.572 | 0.050 | 0.960 | 3.617 | L3Tip |
| **Number of brace roots** | 0.400 | 0.399 | 0.446 | 0.564 | 0.133 | L3Tip |
| **Oil** | 0.340 | 0.323 | 0.287 | 0.723 | 1.716 | L3Tip |
| **Plant height** | 0.341 | 0.264 | 0.030 | 0.980 | 7.644 | L3Tip |
| **Protein** | 0.237 | 0.192 | 0.218 | 0.792 | 4.440 | L3Tip |
| **Southern leaf blight** | 0.402 | 0.369 | 0.129 | 0.881 | 3.292 | L3Tip |
| **Starch** | 0.357 | 0.310 | 0.089 | 0.921 | 4.626 | L3Tip |

| | | | | | | |
|---|---|---|---|---|---|---|
| **Tassel length** | 0.233 | 0.199 | 0.238 | 0.772 | 3.460 | L3Tip |
| **Tassel primary branches** | 0.348 | 0.377 | 0.802 | 0.208 | -2.880 | L3Tip |
| **Total kernel number** | 0.215 | 0.234 | 0.822 | 0.188 | -1.955 | L3Tip |
| **Kernel wt** | 0.353 | 0.347 | 0.337 | 0.673 | 0.642 | L3Tip |
| **Leaf angle** | 0.291 | 0.287 | 0.465 | 0.545 | 0.396 | L3Tip |
| **Test wt** | 0.377 | 0.357 | 0.218 | 0.792 | 1.964 | L3Tip |
| **Cob diameter** | 0.413 | 0.353 | 0.010 | 1.000 | 5.999 | L3Base |
| **Cob length** | 0.136 | 0.155 | 0.752 | 0.257 | -1.901 | L3Base |
| **Cob mass** | 0.289 | 0.274 | 0.317 | 0.693 | 1.536 | L3Base |
| **Ear height** | 0.546 | 0.473 | 0.030 | 0.980 | 7.303 | L3Base |
| **Ear mass** | 0.394 | 0.370 | 0.109 | 0.901 | 2.407 | L3Base |
| **Ear row number** | 0.399 | 0.385 | 0.277 | 0.733 | 1.398 | L3Base |
| **Anthesis silking interval** | 0.010 | 0.053 | 0.851 | 0.158 | -4.321 | L3Base |
| **Days to anthesis** | 0.697 | 0.677 | 0.059 | 0.950 | 2.039 | L3Base |
| **Days to Silking** | 0.656 | 0.644 | 0.218 | 0.792 | 1.185 | L3Base |
| **Kernel number per row** | 0.190 | 0.195 | 0.574 | 0.436 | -0.500 | L3Base |
| **Leaf length** | 0.568 | 0.458 | 0.010 | 1.000 | 11.026 | L3Base |
| **Leaf width** | 0.430 | 0.380 | 0.069 | 0.941 | 4.986 | L3Base |
| **Nodes above ear** | 0.328 | 0.317 | 0.416 | 0.594 | 1.084 | L3Base |
| **Nodes below ear** | 0.599 | 0.576 | 0.119 | 0.891 | 2.361 | L3Base |
| **Number of brace roots** | 0.420 | 0.406 | 0.277 | 0.733 | 1.376 | L3Base |
| **Oil** | 0.310 | 0.316 | 0.545 | 0.465 | -0.629 | L3Base |
| **Plant height** | 0.403 | 0.263 | 0.010 | 1.000 | 14.083 | L3Base |
| **Protein** | 0.223 | 0.188 | 0.188 | 0.822 | 3.536 | L3Base |
| **Southern leaf blight** | 0.405 | 0.369 | 0.089 | 0.921 | 3.590 | L3Base |
| **Starch** | 0.321 | 0.307 | 0.356 | 0.653 | 1.413 | L3Base |

| | | | | | | |
|---|---|---|---|---|---|---|
| **Tassel length** | 0.307 | 0.206 | 0.030 | 0.980 | 10.156 | L3Base |
| **Tassel primary branches** | 0.377 | 0.380 | 0.535 | 0.475 | -0.356 | L3Base |
| **Total kernel number** | 0.254 | 0.240 | 0.277 | 0.733 | 1.441 | L3Base |
| **Kernel wt** | 0.372 | 0.350 | 0.069 | 0.941 | 2.238 | L3Base |
| **Leaf angle** | 0.314 | 0.289 | 0.238 | 0.772 | 2.549 | L3Base |
| **Test wt** | 0.368 | 0.364 | 0.475 | 0.535 | 0.423 | L3Base |
| **Cob diameter** | 0.422 | 0.349 | 0.010 | 1.000 | 7.243 | LMAD |
| **Cob length** | 0.111 | 0.163 | 0.950 | 0.059 | -5.248 | LMAD |
| **Cob mass** | 0.310 | 0.270 | 0.168 | 0.842 | 4.004 | LMAD |
| **Ear height** | 0.554 | 0.480 | 0.020 | 0.990 | 7.419 | LMAD |
| **Ear mass** | 0.387 | 0.367 | 0.139 | 0.871 | 1.991 | LMAD |
| **Ear row number** | 0.423 | 0.378 | 0.069 | 0.941 | 4.564 | LMAD |
| **Anthesis silking interval** | 0.015 | 0.061 | 0.782 | 0.228 | -4.594 | LMAD |
| **Days to anthesis** | 0.737 | 0.677 | 0.010 | 1.000 | 6.014 | LMAD |
| **Days to Silking** | 0.703 | 0.646 | 0.010 | 1.000 | 5.695 | LMAD |
| **Kernel number per row** | 0.167 | 0.196 | 0.812 | 0.198 | -2.887 | LMAD |
| **Leaf length** | 0.534 | 0.453 | 0.010 | 1.000 | 8.148 | LMAD |
| **Leaf width** | 0.420 | 0.383 | 0.119 | 0.891 | 3.713 | LMAD |
| **Nodes above ear** | 0.360 | 0.311 | 0.050 | 0.960 | 4.885 | LMAD |
| **Nodes below ear** | 0.620 | 0.574 | 0.020 | 0.990 | 4.545 | LMAD |
| **Number of brace roots** | 0.435 | 0.405 | 0.158 | 0.851 | 2.975 | LMAD |
| **Oil** | 0.263 | 0.318 | 0.970 | 0.040 | -5.445 | LMAD |
| **Plant height** | 0.329 | 0.277 | 0.089 | 0.921 | 5.263 | LMAD |
| **Protein** | 0.131 | 0.197 | 0.921 | 0.089 | -6.605 | LMAD |
| **Southern leaf blight** | 0.388 | 0.369 | 0.248 | 0.762 | 1.954 | LMAD |
| **Starch** | 0.262 | 0.310 | 0.950 | 0.059 | -4.841 | LMAD |

| | | | | | | |
|---|---|---|---|---|---|---|
| **Tassel length** | 0.291 | 0.205 | 0.050 | 0.960 | 8.622 | LMAD |
| **Tassel primary branches** | 0.353 | 0.383 | 0.772 | 0.238 | -3.000 | LMAD |
| **Total kernel number** | 0.210 | 0.238 | 0.792 | 0.218 | -2.857 | LMAD |
| **Kernel wt** | 0.361 | 0.349 | 0.257 | 0.752 | 1.240 | LMAD |
| **Leaf angle** | 0.334 | 0.286 | 0.040 | 0.970 | 4.785 | LMAD |
| **Test wt** | 0.354 | 0.358 | 0.574 | 0.436 | -0.428 | LMAD |
| **Cob diameter** | 0.357 | 0.355 | 0.475 | 0.535 | 0.181 | LMAN |
| **Cob length** | 0.133 | 0.167 | 0.832 | 0.178 | -3.362 | LMAN |
| **Cob mass** | 0.180 | 0.276 | 1.000 | 0.010 | -9.632 | LMAN |
| **Ear height** | 0.569 | 0.476 | 0.020 | 0.990 | 9.353 | LMAN |
| **Ear mass** | 0.338 | 0.370 | 0.901 | 0.109 | -3.245 | LMAN |
| **Ear row number** | 0.355 | 0.379 | 0.782 | 0.228 | -2.383 | LMAN |
| **Anthesis silking interval** | 0.075 | 0.051 | 0.327 | 0.683 | 2.412 | LMAN |
| **Days to anthesis** | 0.759 | 0.678 | 0.010 | 1.000 | 8.061 | LMAN |
| **Days to Silking** | 0.720 | 0.647 | 0.010 | 1.000 | 7.319 | LMAN |
| **Kernel number per row** | 0.180 | 0.197 | 0.663 | 0.347 | -1.660 | LMAN |
| **Leaf length** | 0.552 | 0.454 | 0.010 | 1.000 | 9.792 | LMAN |
| **Leaf width** | 0.397 | 0.385 | 0.366 | 0.644 | 1.180 | LMAN |
| **Nodes above ear** | 0.377 | 0.315 | 0.020 | 0.990 | 6.207 | LMAN |
| **Nodes below ear** | 0.632 | 0.571 | 0.010 | 1.000 | 6.081 | LMAN |
| **Number of brace roots** | 0.445 | 0.397 | 0.030 | 0.980 | 4.878 | LMAN |
| **Oil** | 0.291 | 0.319 | 0.832 | 0.178 | -2.815 | LMAN |
| **Plant height** | 0.335 | 0.273 | 0.129 | 0.881 | 6.236 | LMAN |
| **Protein** | 0.125 | 0.194 | 0.901 | 0.109 | -6.847 | LMAN |
| **Southern leaf blight** | 0.411 | 0.360 | 0.040 | 0.970 | 5.046 | LMAN |
| **Starch** | 0.314 | 0.311 | 0.485 | 0.525 | 0.315 | LMAN |

| | | | | | | |
|---|---|---|---|---|---|---|
| **Tassel length** | 0.162 | 0.209 | 0.881 | 0.129 | -4.722 | LMAN |
| **Tassel primary branches** | 0.379 | 0.378 | 0.505 | 0.505 | 0.094 | LMAN |
| **Total kernel number** | 0.214 | 0.241 | 0.782 | 0.228 | -2.710 | LMAN |
| **Kernel wt** | 0.330 | 0.351 | 0.851 | 0.158 | -2.135 | LMAN |
| **Leaf angle** | 0.353 | 0.284 | 0.020 | 0.990 | 6.901 | LMAN |
| **Test wt** | 0.326 | 0.364 | 0.911 | 0.099 | -3.792 | LMAN |
| **Cob diameter** | 0.359 | 0.349 | 0.356 | 0.653 | 0.985 | GShoot |
| **Cob length** | 0.160 | 0.168 | 0.614 | 0.396 | -0.764 | GShoot |
| **Cob mass** | 0.249 | 0.277 | 0.822 | 0.188 | -2.848 | GShoot |
| **Ear height** | 0.566 | 0.476 | 0.020 | 0.990 | 8.937 | GShoot |
| **Ear mass** | 0.401 | 0.367 | 0.040 | 0.970 | 3.387 | GShoot |
| **Ear row number** | 0.402 | 0.381 | 0.257 | 0.752 | 2.161 | GShoot |
| **Anthesis silking interval** | 0.050 | 0.058 | 0.614 | 0.396 | -0.829 | GShoot |
| **Days to anthesis** | 0.716 | 0.679 | 0.010 | 1.000 | 3.695 | GShoot |
| **Days to Silking** | 0.687 | 0.648 | 0.010 | 1.000 | 3.942 | GShoot |
| **Kernel number per row** | 0.244 | 0.197 | 0.089 | 0.921 | 4.729 | GShoot |
| **Leaf length** | 0.568 | 0.458 | 0.010 | 1.000 | 10.995 | GShoot |
| **Leaf width** | 0.399 | 0.386 | 0.416 | 0.594 | 1.254 | GShoot |
| **Nodes above ear** | 0.355 | 0.311 | 0.089 | 0.921 | 4.334 | GShoot |
| **Nodes below ear** | 0.623 | 0.573 | 0.010 | 1.000 | 5.033 | GShoot |
| **Number of brace roots** | 0.448 | 0.403 | 0.069 | 0.941 | 4.432 | GShoot |
| **Oil** | 0.360 | 0.310 | 0.059 | 0.950 | 4.961 | GShoot |
| **Plant height** | 0.426 | 0.272 | 0.020 | 0.990 | 15.383 | GShoot |
| **Protein** | 0.305 | 0.186 | 0.010 | 1.000 | 11.957 | GShoot |
| **Southern leaf blight** | 0.407 | 0.371 | 0.059 | 0.950 | 3.643 | GShoot |
| **Starch** | 0.339 | 0.309 | 0.168 | 0.842 | 3.024 | GShoot |

| | | | | | | |
|---|---|---|---|---|---|---|
| **Tassel length** | 0.308 | 0.207 | 0.020 | 0.990 | 10.084 | GShoot |
| **Tassel primary branches** | 0.383 | 0.381 | 0.545 | 0.465 | 0.184 | GShoot |
| **Total kernel number** | 0.303 | 0.236 | 0.020 | 0.990 | 6.687 | GShoot |
| **Kernel wt** | 0.385 | 0.347 | 0.030 | 0.980 | 3.716 | GShoot |
| **Leaf angle** | 0.343 | 0.291 | 0.069 | 0.941 | 5.138 | GShoot |
| **Test wt** | 0.369 | 0.364 | 0.406 | 0.604 | 0.520 | GShoot |
| **Cob diameter** | 0.386 | 0.350 | 0.069 | 0.941 | 3.641 | GRoot |
| **Cob length** | 0.078 | 0.159 | 0.990 | 0.020 | -8.109 | GRoot |
| **Cob mass** | 0.266 | 0.276 | 0.673 | 0.337 | -1.000 | GRoot |
| **Ear height** | 0.604 | 0.474 | 0.010 | 1.000 | 12.951 | GRoot |
| **Ear mass** | 0.365 | 0.369 | 0.634 | 0.376 | -0.369 | GRoot |
| **Ear row number** | 0.335 | 0.379 | 0.941 | 0.069 | -4.348 | GRoot |
| **Anthesis silking interval** | 0.038 | 0.051 | 0.614 | 0.396 | -1.347 | GRoot |
| **Days to anthesis** | 0.752 | 0.675 | 0.010 | 1.000 | 7.670 | GRoot |
| **Days to Silking** | 0.719 | 0.644 | 0.010 | 1.000 | 7.591 | GRoot |
| **Kernel number per row** | 0.174 | 0.197 | 0.792 | 0.218 | -2.318 | GRoot |
| **Leaf length** | 0.565 | 0.453 | 0.010 | 1.000 | 11.190 | GRoot |
| **Leaf width** | 0.390 | 0.378 | 0.455 | 0.554 | 1.235 | GRoot |
| **Nodes above ear** | 0.356 | 0.313 | 0.089 | 0.921 | 4.331 | GRoot |
| **Nodes below ear** | 0.640 | 0.571 | 0.010 | 1.000 | 6.968 | GRoot |
| **Number of brace roots** | 0.428 | 0.400 | 0.178 | 0.832 | 2.714 | GRoot |
| **Oil** | 0.343 | 0.317 | 0.218 | 0.792 | 2.557 | GRoot |
| **Plant height** | 0.380 | 0.268 | 0.020 | 0.990 | 11.174 | GRoot |
| **Protein** | 0.267 | 0.181 | 0.040 | 0.970 | 8.611 | GRoot |
| **Southern leaf blight** | 0.412 | 0.363 | 0.030 | 0.980 | 4.904 | GRoot |
| **Starch** | 0.314 | 0.304 | 0.455 | 0.554 | 1.025 | GRoot |

| | | | | | | |
|---|---|---|---|---|---|---|
| **Tassel length** | 0.277 | 0.202 | 0.079 | 0.931 | 7.556 | GRoot |
| **Tassel primary branches** | 0.371 | 0.378 | 0.545 | 0.465 | -0.739 | GRoot |
| **Total kernel number** | 0.239 | 0.242 | 0.515 | 0.495 | -0.282 | GRoot |
| **Kernel wt** | 0.339 | 0.349 | 0.693 | 0.317 | -1.071 | GRoot |
| **Leaf angle** | 0.273 | 0.279 | 0.574 | 0.436 | -0.570 | GRoot |
| **Test wt** | 0.387 | 0.360 | 0.257 | 0.752 | 2.640 | GRoot |
| **Cob diameter** | 0.379 | 0.353 | 0.178 | 0.832 | 2.590 | Kern |
| **Cob length** | 0.248 | 0.160 | 0.020 | 0.990 | 8.727 | Kern |
| **Cob mass** | 0.276 | 0.268 | 0.416 | 0.594 | 0.770 | Kern |
| **Ear height** | 0.565 | 0.475 | 0.010 | 1.000 | 9.019 | Kern |
| **Ear mass** | 0.403 | 0.369 | 0.040 | 0.970 | 3.396 | Kern |
| **Ear row number** | 0.404 | 0.383 | 0.228 | 0.782 | 2.168 | Kern |
| **Anthesis silking interval** | 0.046 | 0.049 | 0.525 | 0.485 | -0.284 | Kern |
| **Days to anthesis** | 0.728 | 0.677 | 0.010 | 1.000 | 5.176 | Kern |
| **Days to Silking** | 0.687 | 0.645 | 0.020 | 0.990 | 4.200 | Kern |
| **Kernel number per row** | 0.245 | 0.197 | 0.079 | 0.931 | 4.849 | Kern |
| **Leaf length** | 0.549 | 0.458 | 0.010 | 1.000 | 9.084 | Kern |
| **Leaf width** | 0.420 | 0.381 | 0.119 | 0.891 | 3.909 | Kern |
| **Nodes above ear** | 0.365 | 0.311 | 0.079 | 0.931 | 5.469 | Kern |
| **Nodes below ear** | 0.627 | 0.579 | 0.010 | 1.000 | 4.877 | Kern |
| **Number of brace roots** | 0.442 | 0.402 | 0.040 | 0.970 | 3.968 | Kern |
| **Oil** | 0.352 | 0.316 | 0.119 | 0.891 | 3.553 | Kern |
| **Plant height** | 0.407 | 0.263 | 0.010 | 1.000 | 14.360 | Kern |
| **Protein** | 0.166 | 0.177 | 0.604 | 0.406 | -1.064 | Kern |
| **Southern leaf blight** | 0.423 | 0.364 | 0.020 | 0.990 | 5.951 | Kern |
| **Starch** | 0.299 | 0.304 | 0.545 | 0.465 | -0.519 | Kern |

| | | | | | | |
|---|---|---|---|---|---|---|
| **Tassel length** | 0.278 | 0.204 | 0.030 | 0.980 | 7.417 | Kern |
| **Tassel primary branches** | 0.372 | 0.379 | 0.564 | 0.446 | -0.644 | Kern |
| **Total kernel number** | 0.283 | 0.242 | 0.089 | 0.921 | 4.137 | Kern |
| **Kernel wt** | 0.384 | 0.350 | 0.050 | 0.960 | 3.344 | Kern |
| **Leaf angle** | 0.312 | 0.287 | 0.238 | 0.772 | 2.467 | Kern |
| **Test wt** | 0.405 | 0.365 | 0.099 | 0.911 | 3.985 | Kern |
| **Cob diameter** | 0.405 | 0.353 | 0.010 | 1.000 | 5.243 | mean |
| **Cob length** | 0.141 | 0.161 | 0.733 | 0.277 | -2.018 | mean |
| **Cob mass** | 0.291 | 0.274 | 0.297 | 0.713 | 1.692 | mean |
| **Ear height** | 0.563 | 0.481 | 0.010 | 1.000 | 8.230 | mean |
| **Ear mass** | 0.362 | 0.370 | 0.663 | 0.347 | -0.812 | mean |
| **Ear row number** | 0.354 | 0.384 | 0.871 | 0.139 | -3.053 | mean |
| **Anthesis silking interval** | 0.039 | 0.056 | 0.653 | 0.356 | -1.653 | mean |
| **Days to anthesis** | 0.748 | 0.679 | 0.010 | 1.000 | 6.858 | mean |
| **Days to Silking** | 0.716 | 0.648 | 0.010 | 1.000 | 6.797 | mean |
| **Kernel number per row** | 0.164 | 0.200 | 0.901 | 0.109 | -3.517 | mean |
| **Leaf length** | 0.606 | 0.462 | 0.010 | 1.000 | 14.452 | mean |
| **Leaf width** | 0.412 | 0.388 | 0.287 | 0.723 | 2.321 | mean |
| **Nodes above ear** | 0.369 | 0.315 | 0.040 | 0.970 | 5.409 | mean |
| **Nodes below ear** | 0.633 | 0.576 | 0.010 | 1.000 | 5.638 | mean |
| **Number of brace roots** | 0.423 | 0.400 | 0.198 | 0.812 | 2.275 | mean |
| **Oil** | 0.349 | 0.320 | 0.188 | 0.822 | 2.942 | mean |
| **Plant height** | 0.391 | 0.274 | 0.020 | 0.990 | 11.682 | mean |
| **Protein** | 0.271 | 0.195 | 0.040 | 0.970 | 7.610 | mean |
| **Southern leaf blight** | 0.432 | 0.369 | 0.020 | 0.990 | 6.276 | mean |
| **Starch** | 0.293 | 0.309 | 0.703 | 0.307 | -1.600 | mean |

| | | | | | | |
|---|---|---|---|---|---|---|
| **Tassel length** | 0.283 | 0.206 | 0.050 | 0.960 | 7.720 | mean |
| **Tassel primary branches** | 0.394 | 0.380 | 0.356 | 0.653 | 1.407 | mean |
| **Total kernel number** | 0.201 | 0.243 | 0.960 | 0.050 | -4.126 | mean |
| **Kernel wt** | 0.326 | 0.352 | 0.921 | 0.089 | -2.579 | mean |
| **Leaf angle** | 0.324 | 0.292 | 0.129 | 0.881 | 3.274 | mean |
| **Test wt** | 0.378 | 0.362 | 0.337 | 0.673 | 1.536 | mean |
| **Cob diameter** | 0.406 | 0.351 | 0.030 | 0.980 | 5.506 | max |
| **Cob length** | 0.136 | 0.161 | 0.772 | 0.238 | -2.519 | max |
| **Cob mass** | 0.306 | 0.276 | 0.218 | 0.792 | 3.065 | max |
| **Ear height** | 0.590 | 0.481 | 0.010 | 1.000 | 10.980 | max |
| **Ear mass** | 0.377 | 0.371 | 0.376 | 0.634 | 0.606 | max |
| **Ear row number** | 0.377 | 0.386 | 0.693 | 0.317 | -0.937 | max |
| **Anthesis silking interval** | 0.090 | 0.053 | 0.188 | 0.822 | 3.738 | max |
| **Days to anthesis** | 0.755 | 0.679 | 0.010 | 1.000 | 7.588 | max |
| **Days to Silking** | 0.729 | 0.648 | 0.010 | 1.000 | 8.123 | max |
| **Kernel number per row** | 0.180 | 0.201 | 0.782 | 0.228 | -2.056 | max |
| **Leaf length** | 0.614 | 0.463 | 0.010 | 1.000 | 15.098 | max |
| **Leaf width** | 0.402 | 0.386 | 0.297 | 0.713 | 1.518 | max |
| **Nodes above ear** | 0.392 | 0.312 | 0.010 | 1.000 | 7.931 | max |
| **Nodes below ear** | 0.634 | 0.576 | 0.010 | 1.000 | 5.759 | max |
| **Number of brace roots** | 0.393 | 0.403 | 0.683 | 0.327 | -0.998 | max |
| **Oil** | 0.335 | 0.322 | 0.366 | 0.644 | 1.326 | max |
| **Plant height** | 0.442 | 0.273 | 0.010 | 1.000 | 16.985 | max |
| **Protein** | 0.246 | 0.186 | 0.069 | 0.941 | 5.930 | max |
| **Southern leaf blight** | 0.407 | 0.369 | 0.040 | 0.970 | 3.750 | max |
| **Starch** | 0.319 | 0.305 | 0.317 | 0.693 | 1.385 | max |

| | | | | | | |
|---|---|---|---|---|---|---|
| **Tassel length** | 0.266 | 0.210 | 0.149 | 0.861 | 5.685 | max |
| **Tassel primary branches** | 0.393 | 0.378 | 0.327 | 0.683 | 1.494 | max |
| **Total kernel number** | 0.219 | 0.244 | 0.842 | 0.168 | -2.455 | max |
| **Kernel wt** | 0.347 | 0.352 | 0.624 | 0.386 | -0.518 | max |
| **Leaf angle** | 0.313 | 0.289 | 0.218 | 0.792 | 2.329 | max |
| **Test wt** | 0.362 | 0.363 | 0.535 | 0.475 | -0.092 | max |

**S1 Table. Prediction accuracy of 26 complex traits in the Goodman Association panel using HARE from seven diverse tissues: germinating seedlings root (GRoot), germinating seedlings shoot (GShoot), 2 cm from base of leaf 3 (L3Base), two cm from tip of leaf 3 (L3Tip), mature mid-leaf tissue sampled during mid-day (LMAD), mature mid-leaf tissue sampled during mid-night (LMAN), and developing kernels harvested after 350 growing degree days after pollination (Kern), mean, and maximum expression of genes across all tissues. P value (high) and P value (low) were calculated using a Monte Carlo procedure to test if the accuracy using HARE was significantly higher or lower than random HARE. Models were trained in NAM and tested in Goodman Association panel.**

| Trait | Accuracy using HARE | Mean accuracy using random HARE | P value (high) | P value (low) | Accuracy of HARE over random HARE | Tissues |
|---|---|---|---|---|---|---|
| **Cob diameter** | 0.440 | 0.426 | 0.307 | 0.703 | 1.379 | L3Tip |
| **Cob length** | 0.221 | 0.208 | 0.356 | 0.653 | 1.290 | L3Tip |
| **Cob mass** | 0.450 | 0.404 | 0.030 | 0.980 | 4.553 | L3Tip |
| **Ear height** | 0.486 | 0.432 | 0.020 | 0.990 | 5.376 | L3Tip |
| **Ear mass** | 0.449 | 0.425 | 0.129 | 0.881 | 2.403 | L3Tip |
| **Ear row number** | 0.173 | 0.196 | 0.851 | 0.158 | -2.317 | L3Tip |
| **Anthesis silking interval** | 0.061 | 0.054 | 0.406 | 0.604 | 0.724 | L3Tip |
| **Days to anthesis** | 0.751 | 0.742 | 0.257 | 0.752 | 0.943 | L3Tip |
| **Days to Silking** | 0.697 | 0.689 | 0.376 | 0.634 | 0.803 | L3Tip |
| **Kernel number per row** | 0.160 | 0.157 | 0.396 | 0.614 | 0.319 | L3Tip |
| **Leaf length** | 0.338 | 0.328 | 0.337 | 0.673 | 0.964 | L3Tip |
| **Leaf width** | 0.568 | 0.553 | 0.277 | 0.733 | 1.480 | L3Tip |

| | | | | | | |
|---|---|---|---|---|---|---|
| **Nodes above ear** | 0.307 | 0.348 | 0.901 | 0.109 | -4.182 | L3Tip |
| **Nodes below ear** | 0.619 | 0.591 | 0.089 | 0.921 | 2.826 | L3Tip |
| **Number of brace roots** | 0.331 | 0.303 | 0.069 | 0.941 | 2.794 | L3Tip |
| **Oil** | 0.234 | 0.217 | 0.317 | 0.693 | 1.695 | L3Tip |
| **Plant height** | 0.281 | 0.235 | 0.050 | 0.960 | 4.589 | L3Tip |
| **Protein** | 0.190 | 0.156 | 0.139 | 0.871 | 3.461 | L3Tip |
| **Southern leaf blight** | 0.458 | 0.443 | 0.327 | 0.683 | 1.442 | L3Tip |
| **Starch** | 0.290 | 0.264 | 0.198 | 0.812 | 2.588 | L3Tip |
| **Tassel length** | 0.174 | 0.163 | 0.356 | 0.653 | 1.147 | L3Tip |
| **Tassel primary branches** | 0.240 | 0.219 | 0.208 | 0.802 | 2.105 | L3Tip |
| **Total kernel number** | 0.179 | 0.169 | 0.297 | 0.713 | 0.996 | L3Tip |
| **Kernel wt** | 0.397 | 0.388 | 0.337 | 0.673 | 0.971 | L3Tip |
| **Leaf angle** | 0.359 | 0.332 | 0.158 | 0.851 | 2.729 | L3Tip |
| **Test wt** | 0.313 | 0.320 | 0.673 | 0.337 | -0.723 | L3Tip |
| **Cob diameter** | 0.458 | 0.428 | 0.079 | 0.931 | 2.943 | L3Base |
| **Cob length** | 0.252 | 0.206 | 0.099 | 0.911 | 4.559 | L3Base |
| **Cob mass** | 0.455 | 0.406 | 0.030 | 0.980 | 4.864 | L3Base |
| **Ear height** | 0.462 | 0.433 | 0.149 | 0.861 | 2.886 | L3Base |
| **Ear mass** | 0.468 | 0.429 | 0.020 | 0.990 | 3.877 | L3Base |
| **Ear row number** | 0.162 | 0.195 | 0.901 | 0.109 | -3.381 | L3Base |
| **Anthesis silking interval** | 0.043 | 0.050 | 0.614 | 0.396 | -0.684 | L3Base |
| **Days to anthesis** | 0.751 | 0.742 | 0.317 | 0.693 | 0.921 | L3Base |

| | | | | | | |
|---|---|---|---|---|---|---|
| **Days to Silking** | 0.694 | 0.689 | 0.426 | 0.584 | 0.492 | L3Base |
| **Kernel number per row** | 0.149 | 0.158 | 0.743 | 0.267 | -0.954 | L3Base |
| **Leaf length** | 0.361 | 0.329 | 0.069 | 0.941 | 3.205 | L3Base |
| **Leaf width** | 0.564 | 0.554 | 0.356 | 0.653 | 0.949 | L3Base |
| **Nodes above ear** | 0.312 | 0.348 | 0.891 | 0.119 | -3.533 | L3Base |
| **Nodes below ear** | 0.598 | 0.592 | 0.277 | 0.733 | 0.630 | L3Base |
| **Number of brace roots** | 0.324 | 0.305 | 0.178 | 0.832 | 1.917 | L3Base |
| **Oil** | 0.218 | 0.215 | 0.475 | 0.535 | 0.324 | L3Base |
| **Plant height** | 0.263 | 0.235 | 0.168 | 0.842 | 2.852 | L3Base |
| **Protein** | 0.209 | 0.155 | 0.040 | 0.970 | 5.378 | L3Base |
| **Southern leaf blight** | 0.449 | 0.441 | 0.376 | 0.634 | 0.853 | L3Base |
| **Starch** | 0.328 | 0.262 | 0.020 | 0.990 | 6.624 | L3Base |
| **Tassel length** | 0.220 | 0.160 | 0.020 | 0.990 | 5.999 | L3Base |
| **Tassel primary branches** | 0.258 | 0.224 | 0.139 | 0.871 | 3.420 | L3Base |
| **Total kernel number** | 0.163 | 0.170 | 0.673 | 0.337 | -0.731 | L3Base |
| **Kernel wt** | 0.412 | 0.390 | 0.089 | 0.921 | 2.206 | L3Base |
| **Leaf angle** | 0.366 | 0.332 | 0.069 | 0.941 | 3.423 | L3Base |
| **Test wt** | 0.318 | 0.323 | 0.614 | 0.396 | -0.455 | L3Base |
| **Cob diameter** | 0.463 | 0.427 | 0.050 | 0.960 | 3.639 | LMAD |
| **Cob length** | 0.235 | 0.206 | 0.178 | 0.832 | 2.914 | LMAD |
| **Cob mass** | 0.455 | 0.403 | 0.010 | 1.000 | 5.200 | LMAD |
| **Ear height** | 0.482 | 0.431 | 0.020 | 0.990 | 5.171 | LMAD |

| | | | | | | |
|---|---|---|---|---|---|---|
| **Ear mass** | 0.442 | 0.427 | 0.267 | 0.743 | 1.542 | LMAD |
| **Ear row number** | 0.205 | 0.195 | 0.396 | 0.614 | 0.992 | LMAD |
| **Anthesis silking interval** | 0.072 | 0.053 | 0.228 | 0.782 | 1.953 | LMAD |
| **Days to anthesis** | 0.759 | 0.740 | 0.069 | 0.941 | 1.910 | LMAD |
| **Days to Silking** | 0.705 | 0.688 | 0.129 | 0.881 | 1.727 | LMAD |
| **Kernel number per row** | 0.133 | 0.159 | 0.901 | 0.109 | -2.595 | LMAD |
| **Leaf length** | 0.346 | 0.328 | 0.257 | 0.752 | 1.756 | LMAD |
| **Leaf width** | 0.563 | 0.553 | 0.366 | 0.644 | 1.004 | LMAD |
| **Nodes above ear** | 0.343 | 0.347 | 0.564 | 0.446 | -0.400 | LMAD |
| **Nodes below ear** | 0.616 | 0.589 | 0.089 | 0.921 | 2.717 | LMAD |
| **Number of brace roots** | 0.329 | 0.306 | 0.109 | 0.901 | 2.366 | LMAD |
| **Oil** | 0.217 | 0.223 | 0.515 | 0.495 | -0.616 | LMAD |
| **Plant height** | 0.262 | 0.235 | 0.149 | 0.861 | 2.765 | LMAD |
| **Protein** | 0.180 | 0.159 | 0.277 | 0.733 | 2.103 | LMAD |
| **Southern leaf blight** | 0.446 | 0.439 | 0.366 | 0.644 | 0.721 | LMAD |
| **Starch** | 0.285 | 0.266 | 0.277 | 0.733 | 1.928 | LMAD |
| **Tassel length** | 0.210 | 0.160 | 0.069 | 0.941 | 5.017 | LMAD |
| **Tassel primary branches** | 0.244 | 0.224 | 0.228 | 0.782 | 2.016 | LMAD |
| **Total kernel number** | 0.168 | 0.170 | 0.574 | 0.436 | -0.198 | LMAD |
| **Kernel wt** | 0.394 | 0.388 | 0.416 | 0.594 | 0.513 | LMAD |
| **Leaf angle** | 0.367 | 0.332 | 0.119 | 0.891 | 3.515 | LMAD |

| | | | | | | |
|---|---|---|---|---|---|---|
| **Test wt** | 0.349 | 0.319 | 0.079 | 0.931 | 2.985 | LMAD |
| **Cob diameter** | 0.451 | 0.428 | 0.198 | 0.812 | 2.344 | LMAN |
| **Cob length** | 0.213 | 0.202 | 0.396 | 0.614 | 1.140 | LMAN |
| **Cob mass** | 0.443 | 0.405 | 0.059 | 0.950 | 3.763 | LMAN |
| **Ear height** | 0.479 | 0.430 | 0.030 | 0.980 | 4.894 | LMAN |
| **Ear mass** | 0.447 | 0.425 | 0.188 | 0.822 | 2.197 | LMAN |
| **Ear row number** | 0.183 | 0.196 | 0.703 | 0.307 | -1.335 | LMAN |
| **Anthesis silking interval** | 0.053 | 0.051 | 0.426 | 0.584 | 0.210 | LMAN |
| **Days to anthesis** | 0.770 | 0.742 | 0.030 | 0.980 | 2.798 | LMAN |
| **Days to Silking** | 0.717 | 0.689 | 0.059 | 0.950 | 2.821 | LMAN |
| **Kernel number per row** | 0.144 | 0.155 | 0.713 | 0.297 | -1.058 | LMAN |
| **Leaf length** | 0.367 | 0.326 | 0.050 | 0.960 | 4.071 | LMAN |
| **Leaf width** | 0.582 | 0.551 | 0.040 | 0.970 | 3.112 | LMAN |
| **Nodes above ear** | 0.356 | 0.347 | 0.416 | 0.594 | 0.869 | LMAN |
| **Nodes below ear** | 0.620 | 0.591 | 0.069 | 0.941 | 2.889 | LMAN |
| **Number of brace roots** | 0.337 | 0.301 | 0.040 | 0.970 | 3.638 | LMAN |
| **Oil** | 0.225 | 0.217 | 0.386 | 0.624 | 0.897 | LMAN |
| **Plant height** | 0.271 | 0.235 | 0.079 | 0.931 | 3.608 | LMAN |
| **Protein** | 0.197 | 0.159 | 0.129 | 0.881 | 3.766 | LMAN |
| **Southern leaf blight** | 0.472 | 0.439 | 0.129 | 0.881 | 3.334 | LMAN |
| **Starch** | 0.294 | 0.265 | 0.188 | 0.822 | 2.926 | LMAN |
| **Tassel length** | 0.168 | 0.161 | 0.406 | 0.604 | 0.705 | LMAN |

| | | | | | | |
|---|---|---|---|---|---|---|
| **Tassel primary branches** | 0.242 | 0.219 | 0.208 | 0.802 | 2.307 | LMAN |
| **Total kernel number** | 0.180 | 0.167 | 0.267 | 0.743 | 1.257 | LMAN |
| **Kernel wt** | 0.395 | 0.385 | 0.307 | 0.703 | 0.996 | LMAN |
| **Leaf angle** | 0.363 | 0.330 | 0.139 | 0.871 | 3.252 | LMAN |
| **Test wt** | 0.326 | 0.320 | 0.475 | 0.535 | 0.576 | LMAN |
| **Cob diameter** | 0.451 | 0.428 | 0.149 | 0.861 | 2.331 | GShoot |
| **Cob length** | 0.258 | 0.207 | 0.030 | 0.980 | 5.126 | GShoot |
| **Cob mass** | 0.463 | 0.405 | 0.010 | 1.000 | 5.776 | GShoot |
| **Ear height** | 0.454 | 0.430 | 0.158 | 0.851 | 2.408 | GShoot |
| **Ear mass** | 0.464 | 0.429 | 0.030 | 0.980 | 3.502 | GShoot |
| **Ear row number** | 0.188 | 0.199 | 0.663 | 0.347 | -1.125 | GShoot |
| **Anthesis silking interval** | 0.064 | 0.051 | 0.327 | 0.683 | 1.320 | GShoot |
| **Days to anthesis** | 0.766 | 0.743 | 0.069 | 0.941 | 2.284 | GShoot |
| **Days to Silking** | 0.715 | 0.691 | 0.079 | 0.931 | 2.395 | GShoot |
| **Kernel number per row** | 0.148 | 0.159 | 0.772 | 0.238 | -1.085 | GShoot |
| **Leaf length** | 0.348 | 0.327 | 0.178 | 0.832 | 2.172 | GShoot |
| **Leaf width** | 0.571 | 0.553 | 0.317 | 0.693 | 1.746 | GShoot |
| **Nodes above ear** | 0.389 | 0.343 | 0.030 | 0.980 | 4.665 | GShoot |
| **Nodes below ear** | 0.604 | 0.590 | 0.228 | 0.782 | 1.411 | GShoot |
| **Number of brace roots** | 0.336 | 0.307 | 0.059 | 0.950 | 2.949 | GShoot |
| **Oil** | 0.230 | 0.213 | 0.307 | 0.703 | 1.714 | GShoot |
| **Plant height** | 0.257 | 0.232 | 0.198 | 0.812 | 2.413 | GShoot |

| | | | | | | |
|---|---|---|---|---|---|---|
| **Protein** | 0.226 | 0.154 | 0.020 | 0.990 | 7.217 | GShoot |
| **Southern leaf blight** | 0.466 | 0.439 | 0.168 | 0.842 | 2.688 | GShoot |
| **Starch** | 0.311 | 0.263 | 0.079 | 0.931 | 4.809 | GShoot |
| **Tassel length** | 0.209 | 0.156 | 0.030 | 0.980 | 5.315 | GShoot |
| **Tassel primary branches** | 0.279 | 0.225 | 0.030 | 0.980 | 5.463 | GShoot |
| **Total kernel number** | 0.172 | 0.172 | 0.515 | 0.495 | 0.081 | GShoot |
| **Kernel wt** | 0.419 | 0.392 | 0.069 | 0.941 | 2.719 | GShoot |
| **Leaf angle** | 0.325 | 0.331 | 0.594 | 0.416 | -0.595 | GShoot |
| **Test wt** | 0.314 | 0.321 | 0.624 | 0.386 | -0.698 | GShoot |
| **Cob diameter** | 0.469 | 0.426 | 0.030 | 0.980 | 4.316 | GRoot |
| **Cob length** | 0.234 | 0.202 | 0.158 | 0.851 | 3.219 | GRoot |
| **Cob mass** | 0.452 | 0.403 | 0.020 | 0.990 | 4.970 | GRoot |
| **Ear height** | 0.454 | 0.433 | 0.198 | 0.812 | 2.158 | GRoot |
| **Ear mass** | 0.450 | 0.428 | 0.158 | 0.851 | 2.169 | GRoot |
| **Ear row number** | 0.211 | 0.195 | 0.277 | 0.733 | 1.644 | GRoot |
| **Anthesis silking interval** | 0.107 | 0.055 | 0.089 | 0.921 | 5.195 | GRoot |
| **Days to anthesis** | 0.765 | 0.741 | 0.059 | 0.950 | 2.330 | GRoot |
| **Days to Silking** | 0.716 | 0.689 | 0.079 | 0.931 | 2.652 | GRoot |
| **Kernel number per row** | 0.157 | 0.159 | 0.564 | 0.446 | -0.156 | GRoot |
| **Leaf length** | 0.356 | 0.327 | 0.139 | 0.871 | 2.836 | GRoot |
| **Leaf width** | 0.589 | 0.552 | 0.050 | 0.960 | 3.713 | GRoot |
| **Nodes above ear** | 0.334 | 0.344 | 0.634 | 0.376 | -1.014 | GRoot |

| | | | | | | |
|---|---|---|---|---|---|---|
| **Nodes below ear** | 0.602 | 0.594 | 0.297 | 0.713 | 0.768 | GRoot |
| **Number of brace roots** | 0.331 | 0.306 | 0.119 | 0.891 | 2.451 | GRoot |
| **Oil** | 0.207 | 0.213 | 0.624 | 0.386 | -0.620 | GRoot |
| **Plant height** | 0.231 | 0.235 | 0.564 | 0.446 | -0.462 | GRoot |
| **Protein** | 0.174 | 0.153 | 0.218 | 0.792 | 2.123 | GRoot |
| **Southern leaf blight** | 0.485 | 0.439 | 0.030 | 0.980 | 4.594 | GRoot |
| **Starch** | 0.308 | 0.259 | 0.030 | 0.980 | 4.901 | GRoot |
| **Tassel length** | 0.195 | 0.161 | 0.139 | 0.871 | 3.354 | GRoot |
| **Tassel primary branches** | 0.243 | 0.224 | 0.218 | 0.792 | 1.845 | GRoot |
| **Total kernel number** | 0.184 | 0.170 | 0.257 | 0.752 | 1.332 | GRoot |
| **Kernel wt** | 0.415 | 0.388 | 0.089 | 0.921 | 2.703 | GRoot |
| **Leaf angle** | 0.367 | 0.330 | 0.089 | 0.921 | 3.720 | GRoot |
| **Test wt** | 0.327 | 0.319 | 0.337 | 0.673 | 0.792 | GRoot |
| **Cob diameter** | 0.464 | 0.426 | 0.040 | 0.970 | 3.833 | Kern |
| **Cob length** | 0.221 | 0.204 | 0.287 | 0.723 | 1.710 | Kern |
| **Cob mass** | 0.454 | 0.400 | 0.010 | 1.000 | 5.455 | Kern |
| **Ear height** | 0.466 | 0.428 | 0.059 | 0.950 | 3.809 | Kern |
| **Ear mass** | 0.477 | 0.424 | 0.010 | 1.000 | 5.278 | Kern |
| **Ear row number** | 0.220 | 0.193 | 0.218 | 0.792 | 2.636 | Kern |
| **Anthesis silking interval** | 0.045 | 0.056 | 0.713 | 0.297 | -1.061 | Kern |
| **Days to anthesis** | 0.759 | 0.740 | 0.139 | 0.871 | 1.880 | Kern |
| **Days to Silking** | 0.706 | 0.687 | 0.178 | 0.832 | 1.844 | Kern |

| | | | | | | |
|---|---|---|---|---|---|---|
| **Kernel number per row** | 0.158 | 0.156 | 0.436 | 0.574 | 0.197 | Kern |
| **Leaf length** | 0.363 | 0.327 | 0.069 | 0.941 | 3.580 | Kern |
| **Leaf width** | 0.583 | 0.547 | 0.069 | 0.941 | 3.639 | Kern |
| **Nodes above ear** | 0.341 | 0.349 | 0.614 | 0.396 | -0.788 | Kern |
| **Nodes below ear** | 0.611 | 0.591 | 0.178 | 0.832 | 2.040 | Kern |
| **Number of brace roots** | 0.326 | 0.304 | 0.149 | 0.861 | 2.195 | Kern |
| **Oil** | 0.225 | 0.214 | 0.416 | 0.594 | 1.094 | Kern |
| **Plant height** | 0.257 | 0.234 | 0.218 | 0.792 | 2.299 | Kern |
| **Protein** | 0.213 | 0.153 | 0.050 | 0.960 | 6.008 | Kern |
| **Southern leaf blight** | 0.455 | 0.438 | 0.277 | 0.733 | 1.659 | Kern |
| **Starch** | 0.286 | 0.265 | 0.218 | 0.792 | 2.171 | Kern |
| **Tassel length** | 0.180 | 0.163 | 0.307 | 0.703 | 1.625 | Kern |
| **Tassel primary branches** | 0.233 | 0.222 | 0.366 | 0.644 | 1.124 | Kern |
| **Total kernel number** | 0.178 | 0.169 | 0.356 | 0.653 | 0.920 | Kern |
| **Kernel wt** | 0.394 | 0.387 | 0.366 | 0.644 | 0.636 | Kern |
| **Leaf angle** | 0.352 | 0.329 | 0.218 | 0.792 | 2.243 | Kern |
| **Test wt** | 0.313 | 0.322 | 0.713 | 0.297 | -0.974 | Kern |
| **Cob diameter** | 0.451 | 0.431 | 0.139 | 0.871 | 2.020 | mean |
| **Cob length** | 0.233 | 0.204 | 0.198 | 0.812 | 2.923 | mean |
| **Cob mass** | 0.463 | 0.407 | 0.010 | 1.000 | 5.648 | mean |
| **Ear height** | 0.449 | 0.433 | 0.248 | 0.762 | 1.680 | mean |
| **Ear mass** | 0.457 | 0.429 | 0.069 | 0.941 | 2.787 | mean |

| | | | | | | |
|---|---|---|---|---|---|---|
| **Ear row number** | 0.176 | 0.200 | 0.822 | 0.188 | -2.364 | mean |
| **Anthesis silking interval** | 0.082 | 0.052 | 0.149 | 0.861 | 2.944 | mean |
| **Days to anthesis** | 0.787 | 0.745 | 0.010 | 1.000 | 4.231 | mean |
| **Days to Silking** | 0.740 | 0.692 | 0.010 | 1.000 | 4.860 | mean |
| **Kernel number per row** | 0.136 | 0.157 | 0.881 | 0.129 | -2.052 | mean |
| **Leaf length** | 0.380 | 0.329 | 0.020 | 0.990 | 5.079 | mean |
| **Leaf width** | 0.579 | 0.555 | 0.099 | 0.911 | 2.350 | mean |
| **Nodes above ear** | 0.416 | 0.347 | 0.010 | 1.000 | 6.937 | mean |
| **Nodes below ear** | 0.631 | 0.592 | 0.020 | 0.990 | 3.907 | mean |
| **Number of brace roots** | 0.300 | 0.307 | 0.673 | 0.337 | -0.756 | mean |
| **Oil** | 0.230 | 0.216 | 0.307 | 0.703 | 1.440 | mean |
| **Plant height** | 0.238 | 0.235 | 0.455 | 0.554 | 0.311 | mean |
| **Protein** | 0.230 | 0.155 | 0.010 | 1.000 | 7.506 | mean |
| **Southern leaf blight** | 0.448 | 0.442 | 0.386 | 0.624 | 0.536 | mean |
| **Starch** | 0.307 | 0.267 | 0.079 | 0.931 | 4.060 | mean |
| **Tassel length** | 0.204 | 0.157 | 0.040 | 0.970 | 4.649 | mean |
| **Tassel primary branches** | 0.266 | 0.225 | 0.050 | 0.960 | 4.046 | mean |
| **Total kernel number** | 0.147 | 0.170 | 0.901 | 0.109 | -2.217 | mean |
| **Kernel wt** | 0.403 | 0.393 | 0.347 | 0.663 | 1.024 | mean |
| **Leaf angle** | 0.364 | 0.332 | 0.099 | 0.911 | 3.182 | mean |
| **Test wt** | 0.301 | 0.318 | 0.792 | 0.218 | -1.686 | mean |

| | | | | | | |
|---|---|---|---|---|---|---|
| **Cob diameter** | 0.479 | 0.432 | 0.020 | 0.990 | 4.700 | max |
| **Cob length** | 0.251 | 0.205 | 0.059 | 0.950 | 4.566 | max |
| **Cob mass** | 0.472 | 0.407 | 0.010 | 1.000 | 6.537 | max |
| **Ear height** | 0.471 | 0.433 | 0.059 | 0.950 | 3.791 | max |
| **Ear mass** | 0.471 | 0.428 | 0.020 | 0.990 | 4.276 | max |
| **Ear row number** | 0.181 | 0.199 | 0.782 | 0.228 | -1.860 | max |
| **Anthesis silking interval** | 0.103 | 0.052 | 0.069 | 0.941 | 5.044 | max |
| **Days to anthesis** | 0.799 | 0.743 | 0.010 | 1.000 | 5.541 | max |
| **Days to Silking** | 0.749 | 0.691 | 0.010 | 1.000 | 5.779 | max |
| **Kernel number per row** | 0.146 | 0.158 | 0.762 | 0.248 | -1.202 | max |
| **Leaf length** | 0.369 | 0.328 | 0.020 | 0.990 | 4.123 | max |
| **Leaf width** | 0.597 | 0.553 | 0.010 | 1.000 | 4.409 | max |
| **Nodes above ear** | 0.383 | 0.346 | 0.119 | 0.891 | 3.740 | max |
| **Nodes below ear** | 0.636 | 0.593 | 0.010 | 1.000 | 4.315 | max |
| **Number of brace roots** | 0.313 | 0.306 | 0.366 | 0.644 | 0.656 | max |
| **Oil** | 0.208 | 0.216 | 0.584 | 0.426 | -0.860 | max |
| **Plant height** | 0.250 | 0.235 | 0.287 | 0.723 | 1.411 | max |
| **Protein** | 0.229 | 0.154 | 0.010 | 1.000 | 7.581 | max |
| **Southern leaf blight** | 0.477 | 0.443 | 0.069 | 0.941 | 3.333 | max |
| **Starch** | 0.302 | 0.264 | 0.079 | 0.931 | 3.864 | max |
| **Tassel length** | 0.194 | 0.161 | 0.149 | 0.861 | 3.307 | max |
| **Tassel primary branches** | 0.256 | 0.225 | 0.129 | 0.881 | 3.099 | max |

| | | | | | | |
|---|---|---|---|---|---|---|
| **Total kernel number** | 0.155 | 0.170 | 0.832 | 0.178 | -1.576 | max |
| **Kernel wt** | 0.406 | 0.390 | 0.238 | 0.772 | 1.643 | max |
| **Leaf angle** | 0.377 | 0.333 | 0.010 | 1.000 | 4.442 | max |
| **Test wt** | 0.302 | 0.319 | 0.802 | 0.208 | -1.735 | max |

**S2 Table. Prediction accuracy of 26 complex traits in NAM using HARE from 7 diverse tissues: germinating seedlings root (GRoot), germinating seedlings shoot (GShoot), 2 cm from base of leaf 3 (L3Base), 2 cm from tip of leaf 3 (L3Tip), mature mid-leaf tissue sampled during mid-day (LMAD), mature mid-leaf tissue sampled during mid-night (LMAN), and developing kernels harvested after 350 growing degree days after pollination (Kern), mean, and maximum expression of genes across all tissues. P value (high) and P value (low) were calculated using a Monte Carlo procedure to test if the accuracy using HARE was significantly higher or lower than random HARE. Models were trained in Goodman Association panel and tested in NAM.**
